# Systematic mutational mapping reveals optimal amyloid formation for RIPK function

**DOI:** 10.64898/2026.02.19.706818

**Authors:** Mariano Martín, Benedetta Bolognesi

## Abstract

Amyloid formation, typically associated with neurodegeneration, can also serve essential biological functions. RIPK1 and RIPK3 kinases assemble functional amyloids via their RHIM domains to drive necrosome formation and necroptosis. Here, we systematically map the sequence-function relationship of these domains using deep mutagenesis combined with massively parallel assays that directly report on amyloid nucleation and necroptotic signaling. Analysis of ∼3,000 mutations across RIPK1 and RIPK3 reveals that both proteins rely on a conserved aliphatic tetrad to nucleate amyloids, but diverge mechanistically: RIPK3 nucleation is largely driven by this core interface, whereas RIPK1 requires a second aliphatic surface to achieve efficient nucleation. Cross-species comparisons with mutational atlases from mouse RIPKs show conservation of these principles, underscoring their evolutionary selection. Strikingly, functional outcomes depend on finely balanced nucleation: variants that either reduce or enhance amyloid formation impair necroptosis. Consistently, human population genetics indicates that such variants are extremely rare, suggesting evolutionary optimization of the RHIM domain at a “sweet-spot” of amyloid propensity required for signaling. By linking quantitative mutational landscapes to functional amyloid assembly, this work uncovers how evolution has tuned the amyloid propensity of RIPKs to enable robust cell death signaling. These insights not only advance our understanding of amyloid biology but also provide a framework for therapeutic modulation of necroptosis and for engineering synthetic amyloids with tailored activities.

## Introduction

Necroptosis is a genetically regulated lytic and inflammatory form of cell death. Although it can be triggered by different upstream signals, the tumor necrosis factor (TNF)-induced pathway is the best-characterised^1^. This signaling pathway is mediated by three key molecules: the receptor-interacting protein kinases 1 and 3 (RIPK1 and RIPK3) and the downstream effector protein mixed lineage kinase domain-like (MLKL). RIPK1 engages RIPK3 through their RIP homotypic interaction motifs (RHIM; Fig. 1a) to form the necrosome complex that activates MLKL. In both RIPKs the RHIM domains are embedded within large intrinsically disordered regions (Fig. 1b) and drive protein homo– and hetero-oligomerization forming amyloid fibrils that acts as a central scaffold for signal amplification and effector activation. Accordingly, disruption of the RHIM domains severely impairs RIPK1 or RIPK3 interaction and reduces necroptotic cell death^2–4^.

**Figure 1.**
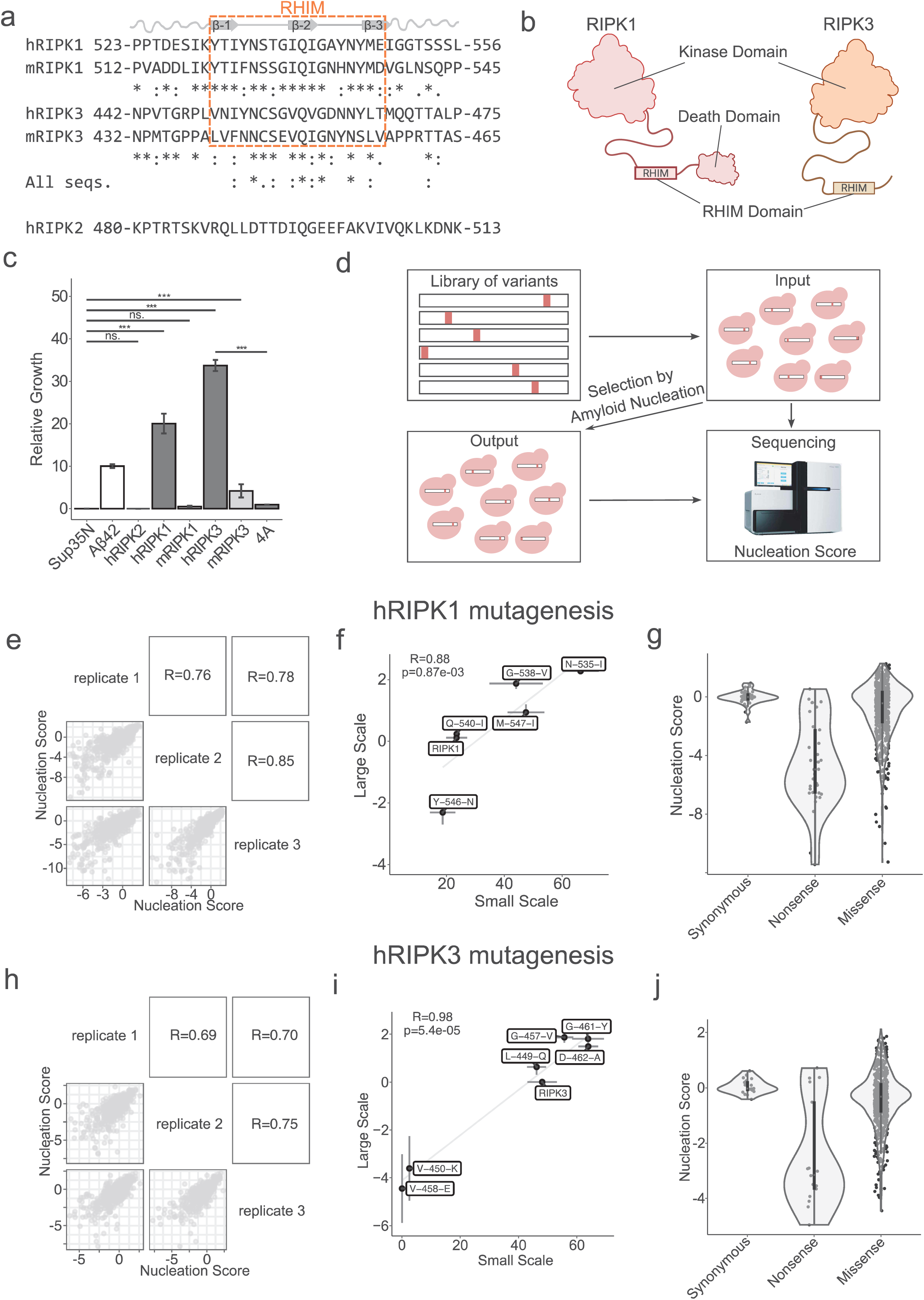
Engineering selection for RIPK1 and RIPK3 amyloid nucleation. **a.** RIPKs sequence alignment. Conservation of residues is represented with ClustalO^43^ consensus symbols: “*” fully conserved residue, “:” conservation of strongly similar amino acids and “.” conservation of weekly similar amino acids. **b.** Schematic cartoon of RIPK1 and RIPK3. **b.** Schematic cartoon of RIPK1 and RIPK3. **c.** Relative growth of cells expressing hRIPK1, hRIPK2, hRIPK3, mRIPK1, mRIPK3 and hRIPK3 VQVG/AAAA (4A) fusions calculated as number of colonies growing in the selective conditions of the amyloid nucleation assay (−URA, −Ade) over colonies growing in the absence of selection (−URA). Error bars indicate SD (n = 3). Cells expressing only Sup35N (no fusion) were used as negative control and cells expressing Sup35N-Aβ42 as positive control. **d.** Deep mutagenesis scheme. A library of sequences of interest is fused to Sup35N and expressed in yeast to be selected for amyloid nucleation. Amyloid nucleation scores are obtained through deep sequencing of input and output samples. **e** and **h.** Correlation of nucleation scores between biological replicates for variants in the hRIPK1 library (e) and hRIPK3 library (h). Pearson’s correlation coefficients are indicated. **f** and **i.** Correlation between nucleation scores obtained from selection and deep sequencing with relative growth of individual variants in selective over nonselective conditions for hRIPK1 (f) and for hRIPK3 (i). Vertical and horizontal error bars indicate estimated sigma errors and SD (n = 3), respectively. Pearson’s correlation coefficients and p-values are indicated. **g** and **j.** Distribution of nucleation scores for each class of mutations in hRIPK1 library (g) and hRIPK3 library (j).

Necroptosis is a central mechanism in tissue homeostasis and in the immune defence against intracellular pathogens. Consistently, mutations in RIPK1 and RIPK3 have been linked to immunologic disorders and autoinflammation^5–8^. Notably, reported mutations typically cause frameshifts leading to early STOP codons or affect the RIPK1 caspase-6 and –8 cleavage site, but none of them directly impact the RHIM domain.

Several viruses and bacteria have developed different strategies to circumvent this inflammatory form of cell death^9,10^. Such is the case of the murine cytomegalovirus (MCMV), the herpes simplex virus 1 (HSV-1) and the Varicella zoster virus (VZV), that encode for the RHIM-containing proteins M45, ICP6 and ORF20, respectively. These viral RHIMs form amyloid structures and interact with RIPK1 and RIPK3 preventing necroptotic cell death^11–13^. RHIM domains are phylogenetically old and consist of 18-22 residues^14^. They have been found across metazoans, with RHIM-like domains also reported in fungi and bacteria^15–17^. Mammalian RHIMs contain a conserved core tetrad (V/I)-Q-(V/I/L/C)-G, which is essential for protein-protein interactions, amyloid assembly and necroptosis^14^.

The homo-amyloid structures of human and mouse RIPK1 and RIPK3 have a similar S-shaped fold formed by three β-strands that form two hydrophobic interfaces, supporting this architecture as a core signaling scaffold^18–22^. A distinct RIPK1:RIPK3 hetero-amyloid arrangement has also been reported, involving two protofibrils assembled through a stacking of alternating monomers that present a serpentine fold^23^. While RIPK1 and RIPK3 interaction is needed for TNF-induced necroptosis, RIPK1-independent mechanisms have also been reported^1^, mainly involving the Z-DNA binding protein 1 (ZBP1) which is also able to form heteromeric assemblies with RIPK3^24^. Regardless of the exact composition of the heteromeric assembly, the signaling pathway converges on the formation of RIPK3 amyloid fibrils and activation of MLKL^25^. Recent work also revealed that the RIPK3 amyloids seeded by RIPK1 fibril fragments adopt a S-shaped fold similar to the one reported for the homo-amyloid structure^22^, further reinforcing the functional relevance of this conserved amyloid architecture in necrosome formation.

Despite the relevance of the RIPK1 and RIPK3 RHIM domains in necroptosis, the functional impact of only a limited number of RHIM mutations have been quantified, mainly in RIPK3. Therefore, how sequence perturbations in RHIM domains affect oligomerization, amyloid fibril formation and necroptosis, and the resulting mechanistic insights remain largely unknown. Herein, we combined deep mutational scanning (DMS) to an aggregation reporter to quantify the effects of mutations on the aggregation of the human RIPK1 and RIPK3 RHIM domains and their mouse orthologous sequences. The resulting mutational atlases reveal a conserved aliphatic tetrad that forms one of the β-strand interfaces present in all the reported RIPK1 and RIPK3 amyloid structures. They also identify gate-keeper residues that constrain amyloid nucleation and reveal a distinct amyloid nucleation architecture in RIPK1 versus RIPK3. Lastly, we find that mutations altering amyloid nucleation and/or fibril stability disrupt necroptosis, supporting a model in which successful necroptosis depends on a finely tuned window of amyloid formation, which is under evolutionary constraint.

## Results

### Quantifying amyloid nucleation of RIPK1 and RIPK3 RHIM domains

We measured the ability of the human and mouse RIPK1 and RIPK3 RHIM domains (Fig. 1a and b) to form amyloid aggregates using a cell-based selection assay where sequences are fused to the nucleation domain of the yeast prion Sup35 (Sup35N) and the aggregation of the fused peptides nucleates the aggregation of endogenous Sup35, a process required for yeast growth in selective conditions (Supp. Fig. 1a)^26,27^. The RHIM domains, flanked by 8 amino acids at the N-terminus and 9 amino acids at the C-terminus were fused to Sup35N (Fig. 1a and b; hRIPK1 523-556, mRIP1 512-545, hRIPK3 442-475 and mRIP3 432-465). As a negative control we cloned a region of human RIPK2 that aligns with the human RIPK1 and RIPK3 RHIM (hRIPK1 and hRIPK3) domains but that does not contain a RHIM domain (region 480-513). Cells expressing sequences bearing the hRIPK1 or hRIPK3 RHIM domains lead to 20.0 ± 2.3 and 33.7 ± 1.3 % of the cells growing in selection media over non-selective conditions, respectively (Fig. 1c). For comparison, this represents a 2– and 3-fold increase compared to the Alzheimer’s amyloid beta peptide Aβ42^27^. In contrast, cells expressing the RIPK2 sequence do not grow in selective conditions, with no difference compared to expression of the reporter without any fusion (Sup35N; ANOVA-Tukey’s post-hoc test). Likewise, cells transformed with the hRIPK3 RHIM mutant VQVG/AAAA (4A), a documented disruptor of RIPK3 amyloid nucleation^4^, grow significantly less than cells expressing the hRIPK3 wild-type (WT) sequence (ANOVA-Tukey’s post-hoc test, Table S1). Finally, cells carrying mouse orthologous sequences grow less than the human RIPKs: 0.5 ± 0.2 and 4.2 ± 1.6 % for mRIPK1 and mRIPK3, respectively; although mRIPK1 sequence do not grow significantly more than Sup35N alone (ANOVA-Tukey’s post-hoc test, Table S1). Taken together these results show that this assay reports on the amyloid aggregation of the RHIM domains.

### A massively parallel assay for quantifying the effect of RIPKs RHIM domain mutations on amyloid nucleation

To investigate and compare the effects of diverse genetic variants on the aggregation of hRIPK1 and hRIPK3 RHIM domains, we measured the impact of all possible amino acid substitutions using the selection assay described above combined with deep sequencing. We designed and synthesized libraries encompassing all single amino acid substitutions from residue 523 to 556 and 442 to 475 of hRIPK1 and hRIPK3, respectively, and expressed them fused to Sup35N in yeast cells that went under selection. We quantified an amyloid nucleation score (NS) for each variant by deep sequencing before and after selection (Fig. 1d)^27^. The resulting NS are reproducible between replicates (Fig. 1e and h) and correlate well with the effects of variants quantified individually (Fig. 1f and i). The effect of mutations also agree with recently reported data showing that the hRIPK1 N545D mutant significantly delays fibril formation^21^. However, likely due to the challenge of purifying monomeric protein, this data is available for less than a handful of RIPK variants.

For both hRIPK1 and hRIPK3, NS distributions revealed strong effects of nonsense mutations towards decreasing amyloid nucleation and a more moderate effect towards reduced nucleation for missense variants (Fig 1. g and j). Altogether, we measured amyloid nucleation for 673 hRIPK1 amino acid substitutions (126 increasing nucleation and 316 reducing, 225 variants with NS similar to WT sequence, Supp. Fig. 1b and e, Z-test, false discovery rate [FDR] = 0.1), and 659 hRIPK3 substitutions (109 variants increasing nucleation and 236 reducing, 243 variants with NS similar to WT sequence, Supp. Fig. 1b and f, Z-test, false discovery rate [FDR] = 0.1). For hRIPK3 we also quantified the effect of 615 single amino acid insertions and 31 deletions, finding that 137 insertions decrease nucleation, 167 increase it and 351 are WT-like (Supp. Fig. 1b and Supp. Fig. 2, Z-test, false discovery rate [FDR] = 0.1). For single amino acid deletions in hRIPK3, we found that 9 decrease nucleation, 6 increase it and 7 are WT-like (Supp. Fig. 1b and Supp. Fig. 2c, Z-test, false discovery rate [FDR] = 0.1). Sequencing cells transformed with hRIPK1 or hRIPK3 libraries before and after induction (no selection) revealed that their growth rates were not confounded by differential toxicity (Supp. Fig. 1d).

**Figure 2.**
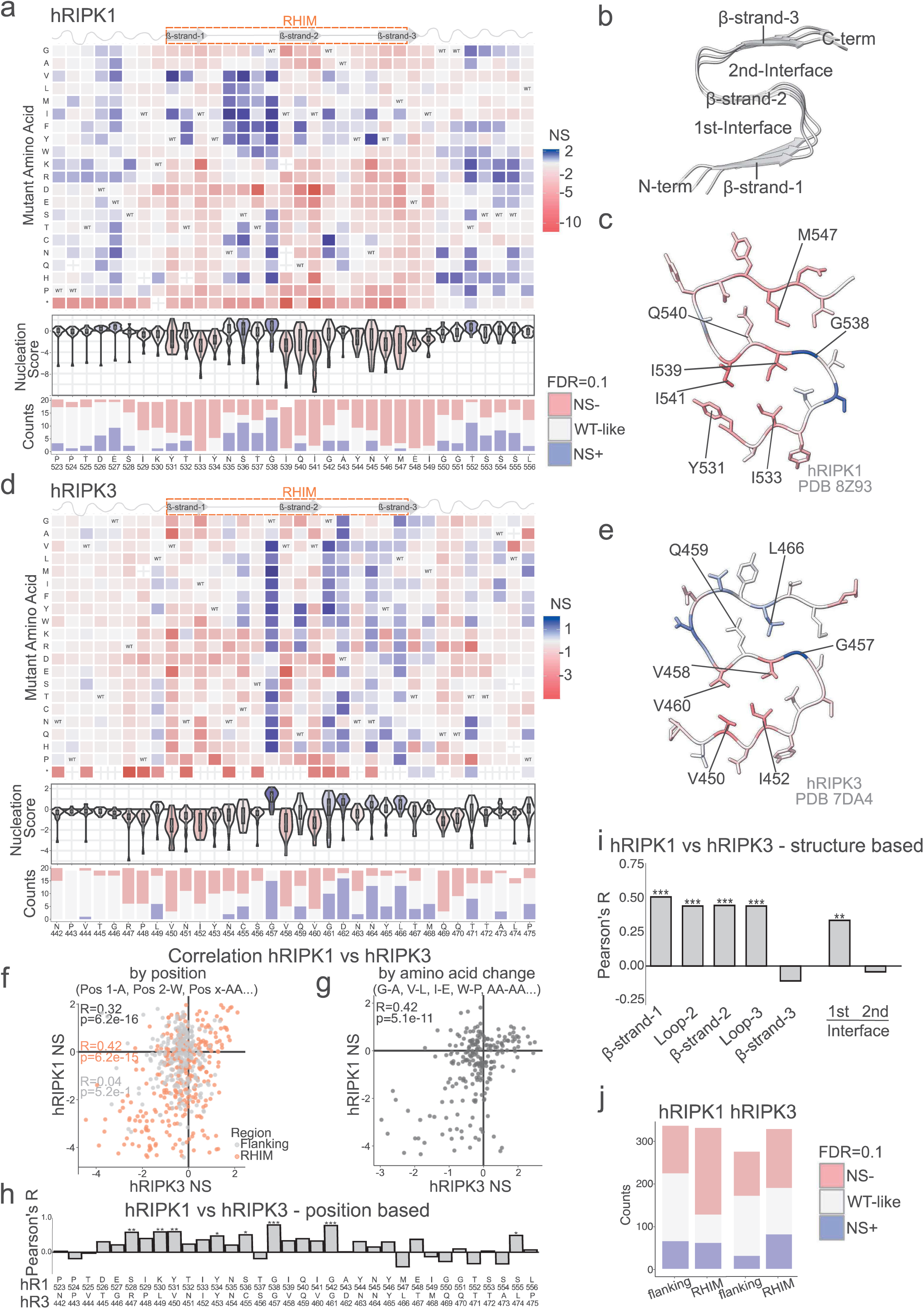
Deep mutagenesis of hRIPK1 and hRIPK3. **a** and **d.** Heatmap of nucleation scores for hRIPK1 (a) and hRIPK3 (d) single amino acid substitutions. The WT amino acid and position are indicated in the x axis and the mutant amino acid is indicated in the y axis. Synonymous variants are indicated as “WT”. The distribution of nucleation scores for each position is summarized in the violin plots below the heatmap. The number of variants increasing or decreasing nucleation at FDR = 0.1 per position are indicated as stacked bars at the bottom. The distribution of nucleation scores for mutations to each amino acid is summarized in the violin plots at the right-hand side of the heatmap. The number of variants increasing or decreasing nucleation at FDR = 0.1 per mutation are indicated as stacked bars at the right. Mutation to stop codons are indicated with an “*”. **b.** Cross-section of the hRIPK1 amyloid fibril (PDB: 9HR6)^21^ indicating the structural elements of the S-shapped fold. **c** and **e.** Monomeric cross-section of the hRIPK1 (c) and hRIPK3 (e) amyloid fibril coloured by the median nucleation score per position. **f.** Correlation of hRIPK1 and hRIPK3 nucleation scores by position. Pearson’s correlation coefficients and p-values are indicated in black for the whole mutagenized sequence, in orange for the RHIM domain and in grey for the flanking regions. **g.** Correlation of hRIPK1 and hRIPK3 nucleation scores by amino acid change. Pearson’s correlation coefficients and p-values are indicated. **h.** Position-based correlation of hRIPK1 and hRIPK3 nucleation scores. Pearson’s correlation coefficient is indicated in the y-axis. p-value: *<0.05, **<0.01 and ***<0.001. **i.** Correlation of hRIPK1 and hRIPK3 nucleation scores for different structural elements from the S-shaped fibril fold. Pearson’s correlation coefficient is indicated in the y-axis. p-value: *<0.05, **<0.01 and ***<0.001. **j.** Number of variants increasing or decreasing nucleation at FDR = 0.1 and variants that are WT-like in the hRIPK1 and hRIPK3 RHIM domains and flanking regions.

### Mutations inside the RHIM domains have more disruptive effects than those in the flanking regions

Inspecting the heatmap of mutational effects for amino acid changes in hRIPK1 and hRIPK3 reveals that most of the mutations that affect amyloid nucleation, either increasing or decreasing it, are localized within the RHIM domains (Fig. 2a and d, Supp. Fig. 1e-f, Chi-squared test RIPK1 p-value=1.398e-13, RIPK3 p-value=1.497e-05). The fact that substitutions to prolines in the RHIM domain all decrease nucleation also suggest that these regions are structured upon nucleation. In contrast, substitutions to prolines are more tolerated in the flanking regions. These results agree with the reported amyloid structures for both hRIPK1 and hRIPK3 that feature a S-shaped structured RHIM domain with disordered or flexible flanking regions at its N-and C-terminus (Fig. 2b, c and e; Supp. Fig. 3; PDB: 8Z93, 9HR9, 9HR6, 7DAC, 7DA4, and 8Z94)^20–22^.

**Figure 3.**
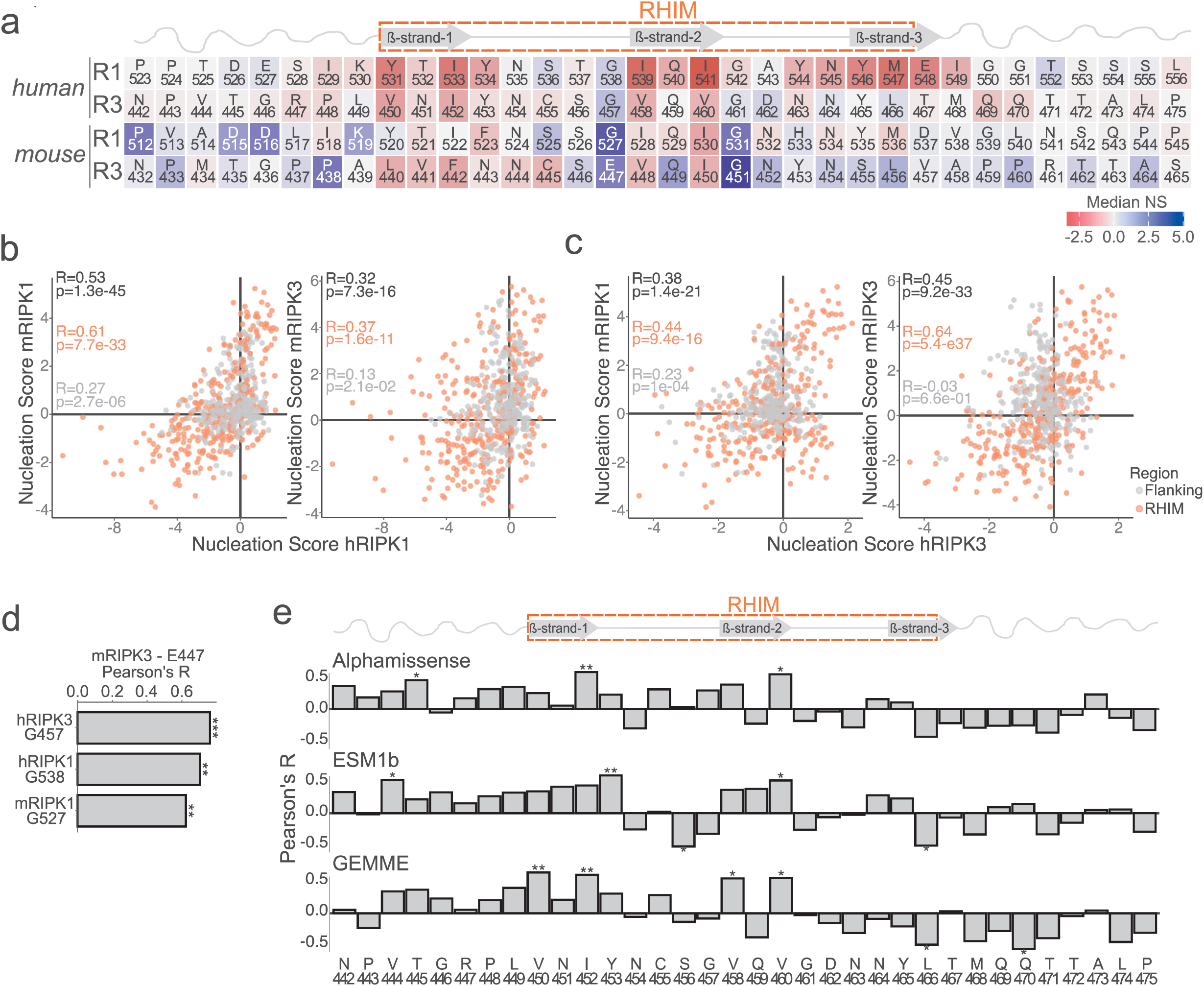
Mutational effects at residues essential for RHIM amyloid nucleation are evolutionarily conserved. **a.** Comparison of median nucleation scores per position across human and mouse RIPK1 and RIPK3 mutational datasets. **b-c.** Correlation of human RIPK1 (b) or RIPK3 (c) with mRIPK1 (left panel) or mRIPK3 (right panel) nucleation scores. Pearson’s correlation coefficients and p-values are indicated in black for the whole mutagenized sequence, in orange for the RHIM domain and in grey for the flanking regions. **d.** Correlation of nucleation scores resulting from mRIPK3 E447 mutations versus mutations of hRIPK3 G457, hRIPK1 G538 and mRIPK1 G527. Pearson’s correlation coefficient is indicated in the x-axis. p-value: *<0.05, **<0.01 and ***<0.001. **e.** Position-based correlation of hRIPK3 nucleation scores versus Alphamissense^28^ (top), ESM1b^29^ (middle) and GEMME^30^ (bottom) scores. Pearson’s correlation coefficient is indicated in the y-axis. p-value: *<0.05, **<0.01 and ***<0.001.

NS correlations show that many of the amino acid changes have similar impact in both sequences (Fig. 2f and g; R=0.32, p=6.2e-16 for residue amino acid change and R=0.42, p=5.1e-11 for type of amino acid change), especially for residues in the RHIM domain (Fig. 2f; R=0.42, p=6.2e-15). In RHIM domains, both sequences present a similar number of mutations that increase nucleation (61 and 79, for hRIPK1 and hRIPK3, respectively) while the main difference stands at the number of variants that decrease nucleation: 204 for hRIPK1 and 135 for hRIPK3 (Fig. 2j). NS correlations per position reveal that most of the significant correlations occur within the RHIM domain (Fig. 2f and h). Positions G538/G457 (hRIPK1/hRIPK3) and G542/G461 present the highest Pearson’s correlation coefficients (Fig. 2h; R=0.87, p-value=1.51e-06 and R=0.85, p-value=3.86e-06), highlighting the relevance of these glycines in the nucleation process of both sequences.

Mutations outside the RHIM show no correlation between the hRIPK1 and hRIPK3 datasets (Fig. 2f; R=0.04, p=5.2e-1) revealing that the flanking regions of both sequences may influence amyloid formation in a distinct way. In particular, in the case of RIPK1 we found that 65 mutations in the flanking regions increase nucleation while for RIPK3 there are only 30 (Fig. 2j). Many mutations at two positions in the RHIM flanking region of hRIPK1, D526 and E527, increase nucleation (6 and 9/19 respectively, Z-test, false discovery rate [FDR] = 0.1; Fig. 2a and Supp. Fig. 1e). These positions act as gatekeepers that prevent excessive amyloid formation. Substitutions to positive charges (R, K and H) outside the RHIM domain also increase nucleation (Fig. 2a and Supp. Fig. 1e). On the other hand, hRIPK3 flanking regions tolerate better substitutions as many of the mutations present minor effects on increasing nucleation propensity (Fig. 2d and Supp. Fig. 1f).

### An aliphatic tetrad is essential for RIPK1 and RIPK3 amyloid nucleation

The reported hRIPK1 and hRIPK3 amyloid structures consist of a S-shaped fold with three β-strands and two hydrophobic interfaces (Fig. 2b). Previous works reported a conserved motif (539-IQIG-542 and 458-VQVG-461 for hRIPK1 and hRIPK3, respectively)^2,4^ which include the central β-strand in these amyloid structures, with residues engaging in both the first (I539 and I541 in hRIPK1; V458 and V460 in hRIPK3) and second hydrophobic interface (Q540 in hRIPK1; Q539 in hRIPK3).

Mapping the median NS per position on the cross-section of the hRIPK1 and hRIPK3 amyloid fibrils (Fig. 2c and e), we find that mutations at residues Y531/V450, I533/I452, I539/V458 and I541/V460 (hRIPK1/hRIPK3 respectively) strongly decrease amyloid nucleation. These residues have the lowest median NS per position and correspond to an aliphatic tetrad conserved in both kinases (Fig. 1a and Supp. Fig. 1c) which forms one of the two hydrophobic interfaces in the amyloid fold (Fig. 2c and e, Sup. Fig. 1e and f). The only other position with similarly low median NS is M547, which is part of the second aliphatic interface in hRIPK1.

Our mutational atlas also reveals that two interfaces are critical for nucleation of hRIPK1, i.e. most mutations decrease NS in these regions (Fig. 2a and c). On the other hand, for hRIPK3, our data prioritizes only the involvement of the first interface in amyloid nucleation (Fig. 2d and e). We further observe a correlation for the effect of mutations in hRIPK1 and hRIPK3 for residues in the first β-strand and loop-2 (Fig. 2i), which is then lost for residues in the third β-strand. In line with this, the effects of mutations affecting the first interface correlate, but not those in the second one (Fig. 2i). Taken together, these results reveal that the aliphatic tetrad forming the first interface is essential for the amyloid nucleation of hRIPK1 and hRIPK3 RHIM domains, while the second interface seems to be relevant only for hRIPK1 amyloid nucleation.

For hRIPK3, we further investigated mutational impact by employing indel mutagenesis, quantifying the effect of 615 single amino acid insertions and 31 single amino acid deletions. Most of the insertions in the first β-strand decrease amyloid nucleation (Supp. Fig. 2a and b), highlighting the relevance of this structural element in the nucleation of the S-shaped amyloid aggregate. In contrast with this, insertion of aliphatic or polar residues before or at the beginning of the second β-strand increases nucleation and almost all insertions affecting the third β-strand increase nucleation. On the other hand, deletions of the tetrad residues I452, V458, Q459 or V460, severely decrease nucleation (Supp. Fig. 2c), revealing the importance of the aliphatic tetrad in the nucleation process. Deletion of specific residues inside loops, such as S456, G457, G461, D462 or N463, increase nucleation.

### Mutations of glycine residues before and after the second β-strand increase nucleation

In both hRIPK1 and hRIPK3, glycine residues flank the second β-strand, at the center of the S-shaped fold of fibrils. Many mutations at these residues increase nucleation, suggesting they may act as gatekeepers of nucleation. This is particularly striking for the first of these glycines (G538 and G457, in gRIPK1 and gRIPK3, respectively), where most of the mutations increase amyloid nucleation (13/19 and 15/19, respectively; Fig. 2c and e, Sup. Fig. 1e-f). In these positions, the effects of mutations in gRIPK1 and gRIPK3 highly correlate (Pearson’s correlation coefficients 0.87 for G538/G457 and 0.85 for G461/G452; Fig. 2f).

### The effect of mutations in residues essential for RHIM amyloid nucleation are evolutionary conserved

RHIM sequences are highly conserved (Fig.1a and Supp. Fig. 1c) but show differing amyloid nucleation propensities, with human RIPKs nucleating more strongly than mouse orthologs (Fig. 1c). To understand these differences, we mutagenized the corresponding mouse regions (mRIPK1 512-545 and mRIPK3 432-465). The obtained nucleation scores are reproducible (Supp. Fig. 4a and e). The effects of mutations in mRIPK3 also agree well with the *in vitro* behaviour of three recombinant mouse variants for which aggregation was measured in the presence of Thioflavin-T^19^ (Supp. Fig. 9g). Overall, we quantified amyloid nucleation for 657 mRIPK1 substitutions (265 increasing nucleation, 200 reducing, 187 similar to WT) and 677 mRIPK3 substitutions (300 increasing, 152 reducing, 208 similar to WT) (Supp. Fig. 4i-l, Z-test, FDR = 0.1), revealing NS distributions for missense mutations which are skewed towards stronger nucleators compared to the human libraries (Supp. Fig. 4b and f).

**Figure 4.**
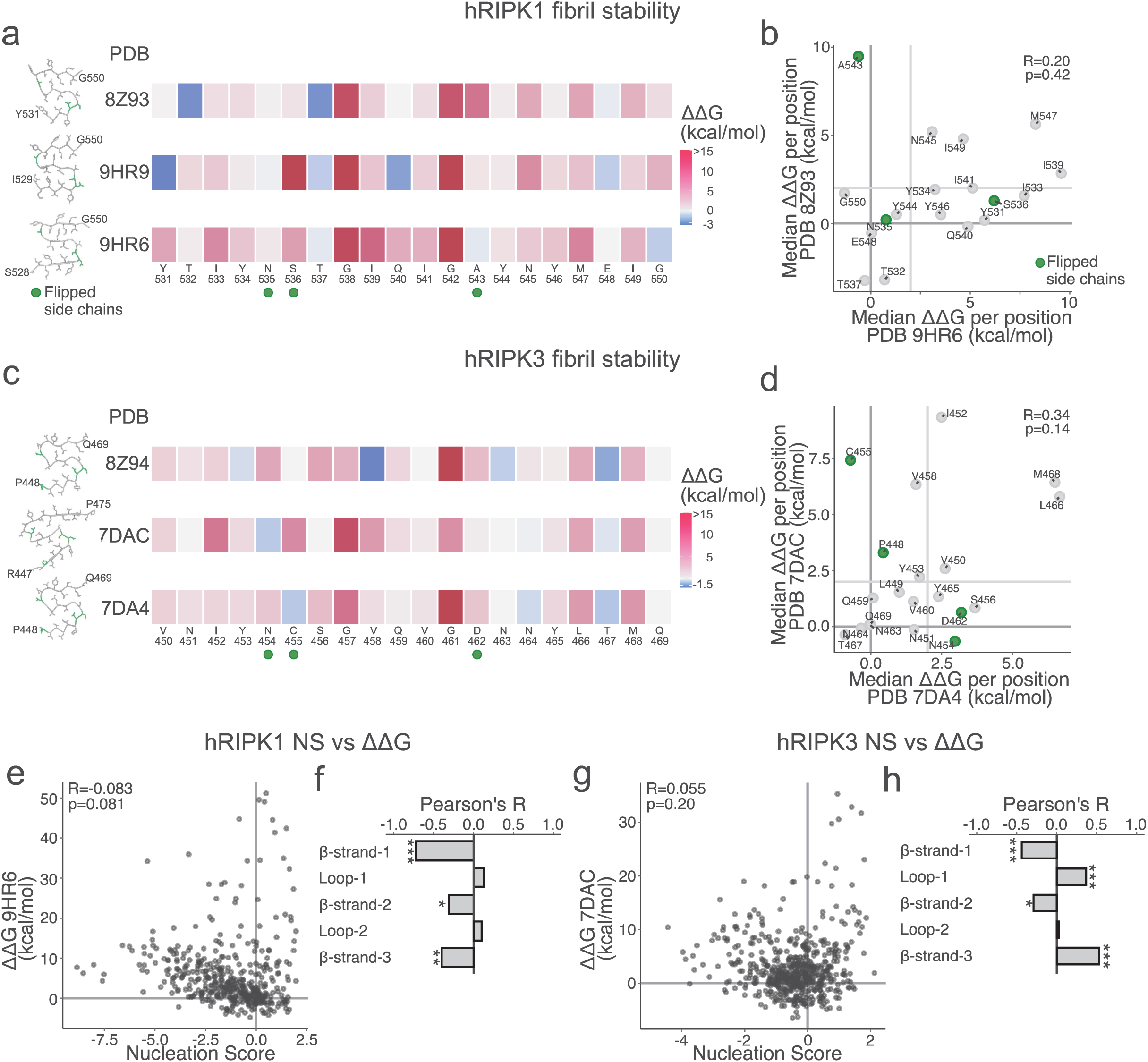
Effect of mutations on hRIPK1 and hRIPK3 fibrils stability. **a** and **c.** Heatmap comparing the median ΔΔG resulting from mutations at each position estimated by FoldX^31^ for different hRIPK1 (a) and hRIPK3 (c) fibrillar structures. Green dots identify those positions where the WT side chains display different orientation amongst PDB structures. **b** and **d.** Correlation of FoldX estimated median ΔΔG per position for different hRIPK1 (b) and hRIPK3 (d) structures. Dashed vertical lines indicates the 2 kcal/mol threshold. **e** and **g.** Correlation between hRIPK1 (e) and hRIPK3 (g) estimated ΔΔGs and nucleation scores per position. **f** and **h.** Nucleation scores and ΔΔG correlations for hRIPK1 (f) and hRIPK3 (h) structural elements present in the S-shaped fold. Pearson’s correlation coefficients and p-values are indicated.

Human and mouse RIPK1 share 55.88% overall identity, 76.47% in the RHIM, similarity values go up to 94.11%; Human and mouse RIPK3 share 41.18% overall identity, 47.06% in the RHIM, with similarity up to 70.59%. In both cases, the most divergent regions are the C-terminal flanking regions (Fig. 1a). We compared median positional effects between sequences (Fig. 3a-c). Orthologs behave similarly (RIPK1: R=0.53, p=1.3e-45; RIPK3: R=0.45, p=9.22e-33), with stronger correlations for mutations inside the RHIM residues (RIPK1: R=0.61, p=7.7e-33; RIPK3: R=0.64, p=5.4e-37) and weaker ones in flanks (RIPK1: R=0.27, p=2.7e-6; RIPK3: R=0.03, p=6.6e-1) (Fig. 3b and c). Cross-ortholog comparisons are weaker but still detectable, especially in RHIM regions (hRIPK1 vs mRIPK3: R=0.37, p=1.6e-11; hRIPK3 vs mRIPK1: R=0.44, p=9.4e-16; Fig. 3b and c).

Between mouse sequences, we observe moderate correlation in RHIM residues (R=0.54, p=1.4e-24) but not in flanking regions (R=0.04, p=5e-01) (Supp. Fig. 4m). Mutation-type analyses also show moderate correlation (R=0.53, p=3.1e-19, Supp. Fig. 4n). As in our observations for the human sequences, major differences arise from mutations impacting the second hydrophobic interface (Supp. Fig. 4j and l).

While hRIPK1, hRIPK3 and mRIPK1 feature a glycine before the second β-strand, the mRIPK3 sequence contains instead glutamic acid. Given the relevance of these residues in the nucleation process, we compared NS for E447 mutations in mRIPK3 with NS for the corresponding glycine mutations in the other sequences and found them highly correlated (Pearssn’s R; mRIPK3 vs hRIPK3: R=0.76, p=6.1e-4; mRIPK3 vs hRIPK1: R=0.70, p=2.3e-3; mRIPK3 vs mRIPK1: R=0.62, p=0.01; Fig. 3d).

Finally, the first two residues of the mRIPK1 aliphatic tetrad (Y520 and I522) appear more mutation-tolerant than their hRIPK1 counterparts (Fig. 3a), but the lower growth rate of the mRIPK1 library may reduce apparent effect sizes.

Overall, RHIM domains are highly conserved and respond similarly to mutations, whereas the less conserved flanking regions behave more divergently across sequences.

### Variant Effect Predictors capture the impact of mutations in RHIM domains at some specific positions

Given the degree of conservation of the mutagenized sequences, we next compared the performance of three Variant Effect Predictors (VEPs), Alphamissense^28^, ESM-1b^29^ and GEMME^30^, in predicting the impact of mutations and their correlation with our datasets. All VEPs scores lack granularity when predicting mutational impact in either hRIPK1 or hRIPK3 RHIM domains. They also do not correlate with NS (Supp. Fig. 5a-c and f-h). However, by evaluating the correlation between VEPs scores and NS position wise, we found a positive correlation between GEMME score and our NS at those positions forming the aliphatic tetrad of hRIPK3, a finding which is not reproduced by Alphamissense and ESM-1b (Fig. 3e). In the case of hRIPK1, when analysing the correlation of VEPs scores and NS position-wise, we found that Alphamissense is the only VEP that correlates with NS in those residues forming the aliphatic tetrad (Supp. Fig. 5e). All of these scores capture the importance of the glycine that is after the second β-strand, G542 and G461 for hRIPK1 and hRIPK3, respectively (Supp. Fig. 5d and i). Despite the VEPs scores at these residues being highly detrimental, they are also highly compressed making a correlation comparison versus the NS not meaningful.

**Figure 5.**
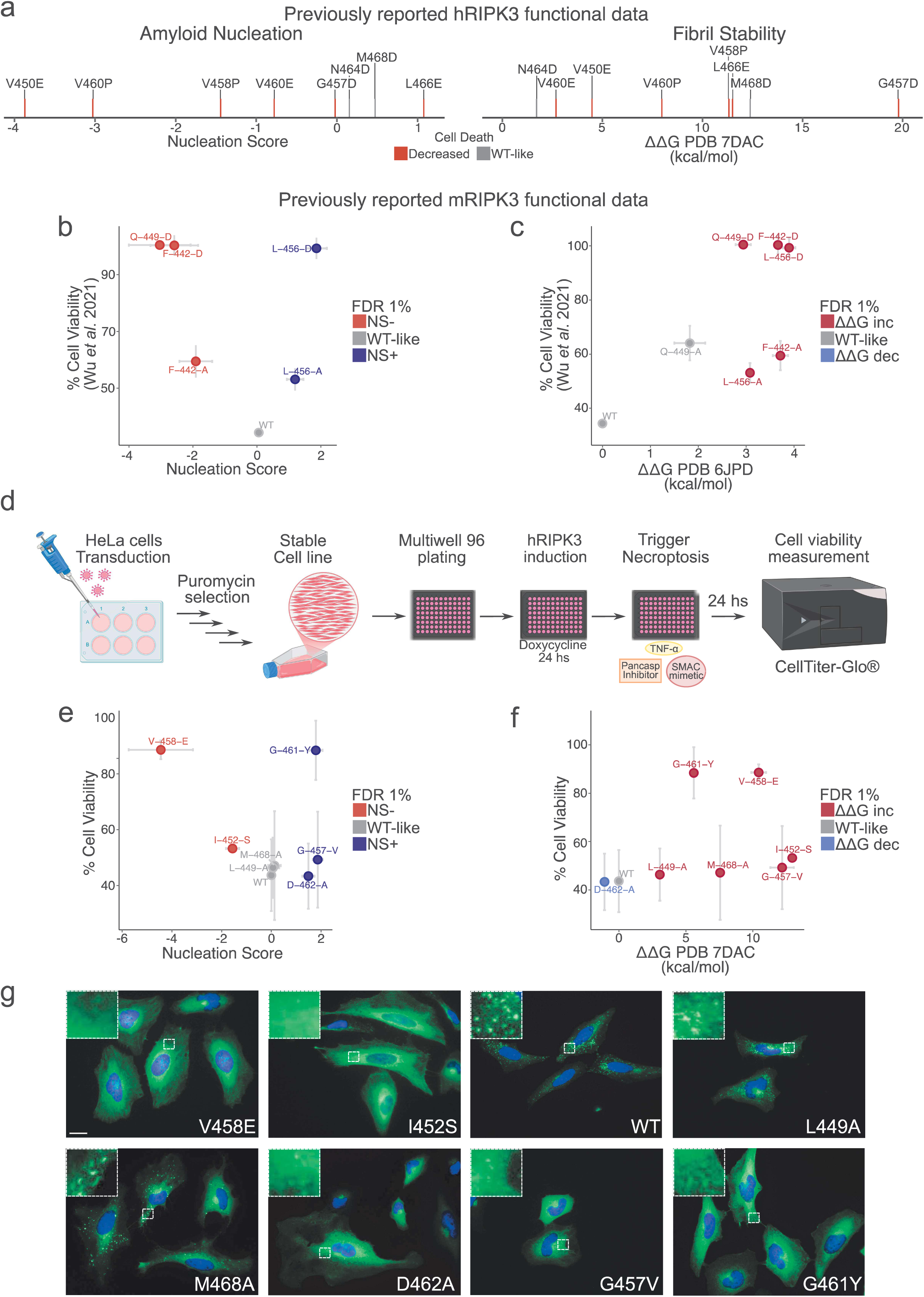
Variants altering RIPK3 amyloid formation disrupt necroptosis. **a.** hRIPK3 variants reported in the literature were classified as decreasing cell death (red) or having a wt-like behavior (grey) and mapped in comparison to hRIPK3 nucleation scores (left) and fibril stability (right). **b-c.** Correlation between previously reported cell survival measurements^19^ and nucleation scores (b) or ΔΔG (c) for mRIPK3 variants. Vertical and horizontal error bars indicate estimated SD (n=3) and sigma errors of the experiments (in the case of the nucleation score) or SD from five FoldX runs (in the case of fibril stability), respectively. **d.** Schematics of the generation of stable HeLa cell lines expressing hRIPK3 or hRIPK3 variants under doxycycline inducible promoter used for quantifying cell viability using CellTiter-Glo® assay. **e-f.** Correlation between cell viability measurements from this study and nucleation scores (e) or FoldX ΔΔGs (f) for hRIPK3 variants. Vertical and horizontal error bars indicate estimated SD (n=3) and sigma errors (in the case of the nucleation score) or SD (in the case of fibril stability) of the experiments, respectively. **g.** Representative immunofluorescence images showing hRIPK3 expression (green) in cells after 4 hs of triggering necroptosis. The nuclei are stained with Dapi (blue). Scale bar 20 μm.

### Mutating aliphatic interfaces or glycines destabilize the amyloid fibrils

We next predicted the impact of mutations on fibril stability by measuring *in silico* the ΔΔG of all variants running FoldX^31^ on all the homo-amyloid structures reported in the human and mouse RIPKs sequences (PDB: 9HR6, 9HR9, 8Z93, 8Z94, 7DAC, 7DA4, 6JPD and 8IB0, Fig. 4a and c and Supp. Fig. 6c, f, g and h)^18–22^. One feature that is highly conserved across the different structures is the destabilizing effect resulting from mutating glycine residues located just before and just after the second β-strand (Fig. 4a and c, Supp. Fig. 6c and f). Glycines are important for allowing turns in the structures and it is likely that any other residues at these positions would be incompatible with the final amyloid fold in these structures. Another common feature across structures is the destabilizing effect of mutating residues forming the hydrophobic interfaces. As expected, mutating residues whose sidechains are exposed to the surface is overall better tolerated (Fig. 4a and c, Supp. Fig. 6c and f).

**Figure 6.**
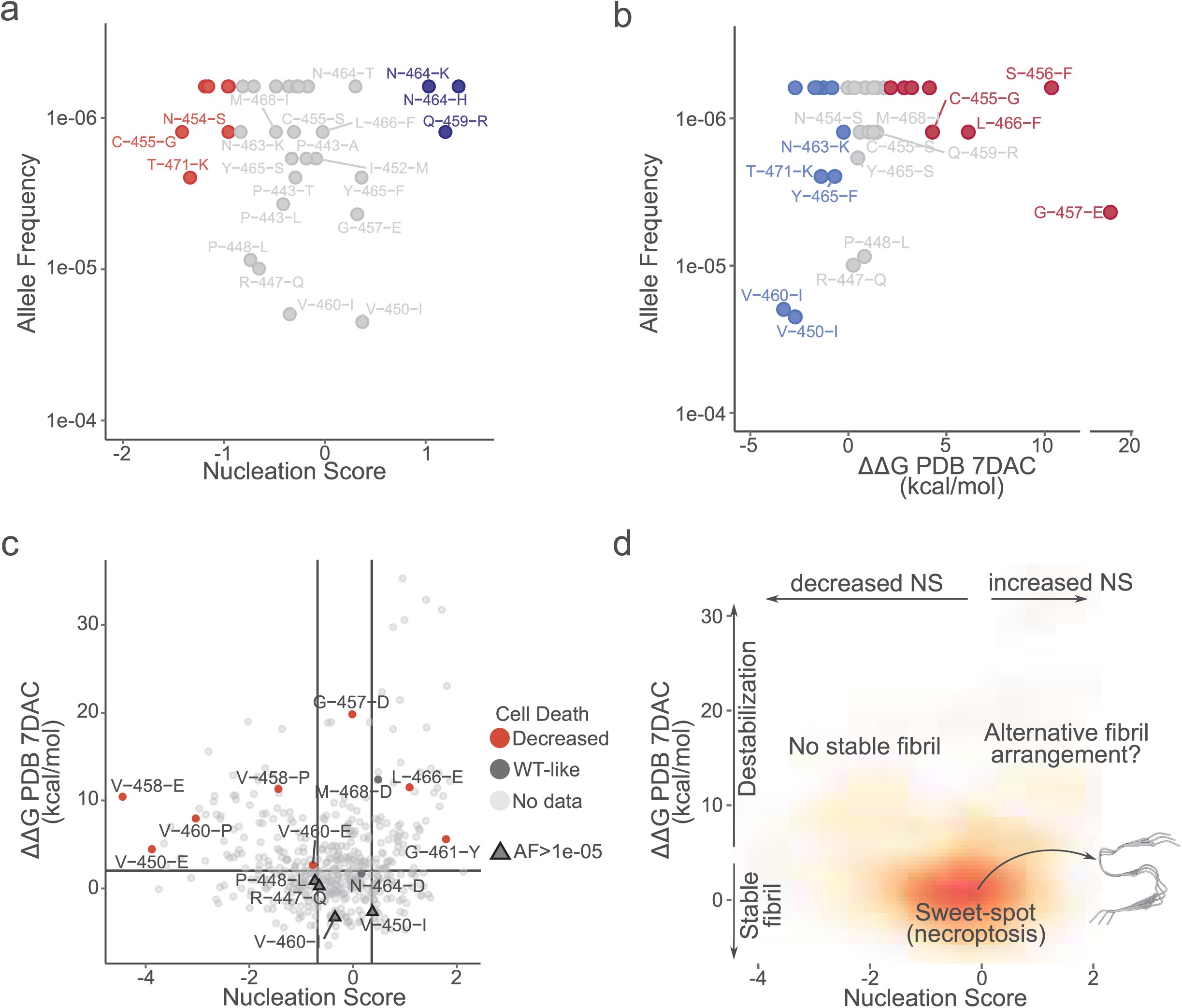
RIPK3 function requires an optimal range of amyloid nucleation and fibril stability. **a-b.** hRIPK3 variants Gnomad^37^ Allele Frequency versus nucleation scores (g) and ΔΔG (h). **c.** Scatter plot of hRIPK3 mutations ΔΔG and nucleation scores. Variants previously studied in the literature, more frequent in the human population (Gnomad Allele Frequency>1e-05) and functionally characterized in this work are highlighted. The horizontal grey line indicates the 2 kcal/mol threshold. Vertical grey lines indicate the maximum and minimum NS of variants decreasing and increasing nucleation at FDR1%, respectively. **d.** Schematics of the “sweet-spot” of amyloid formation needed for necroptosis. The colour scale indicates the density of the variants in each part of the plot. The hRIPK3 fibril structure PDB 7DAC^20^ was used for the FoldX calculations.

For those sequences that have more than one structure reported (hRIPK1 and hRIPK3), we compared the ΔΔG measured in the different structures finding that the effect of mutations on fibril stability is correlated for all of them, except for those residues that feature different side chain orientations (buried vs exposed) (Supp. Fig. 6 and Fig. 4a-d).

### Nucleation Scores correlate with ΔΔG only in β-strands

We next assessed how the NS relates to fibril stability. Across the full dataset, we observed overall no correlation between NS and ΔΔG (Fig. 4e and g, Supp. Fig. 7a, c, e, and g), indicating that our reporter assay does not capture the stability of the resulting amyloid fibrils.

We then examined NS versus ΔΔG within the specific structural elements of the S-shaped amyloid fold. Using structure 9HR6, NS negatively correlate with ΔΔG across all three β-strands of hRIPK1 (Fig. 4f). For hRIPK3 (structure 7DAC), NS shows a negative correlation in the first and second β-strands, but a positive correlation in the first loop and in the β-strand (Fig. 4h). These trends are broadly conserved across alternative structures of the same sequences (Supp. Fig. 7b, d, f, and h).

### RHIM domain mutations that disrupt amyloid formation also impair necroptosis

RHIM domains oligomerize RIPKs through the formation of amyloid fibrils. These structures constitute the cornerstone of the necrosome complex and are central to necroptosis execution. Mutations in the RHIM domain affect this pathway, altering necroptotic cell death^4,19,22,32^. Although the relevance of the amyloid structure for RIPKs function and necroptosis has been reported^4^, the molecular interplay between amyloid formation and kinase activity is still poorly understood. We correlated the reported effect of RIPKs mutations on necroptosis with our NS as well as with ΔΔG estimates from FoldX^31^. For hRIPK3 the effect of eight different RHIM domain singles amino acid substitutions on necroptotic cell death have been previously measured ^4,22,32^. Most of these mutations decrease cell death, meaning that they are disrupting necroptosis, while only two exhibiting WT-like behaviour (Fig. 5a). With the exception of G457D, all mutations that decrease cell death also alter amyloid nucleation, while the two variants that exhibit WT-like function have NS close to WT (Fig. 5a and Supp. Fig. 8a). In terms of stability, most of the mutations that decrease function also decrease fibril stability, while variant M468D maintains WT-like function but is predicted to cause fibril destabilization (Fig. 5a and Supp. Fig 8b-d).

When it comes to mRIPK3 mutations Q449D, F442D and F442A^19^ abrogate necroptosis as well as amyloid nucleation (Fig. 5b). On the other hand, L456A and L456D also decrease function but exhibit an increased NS in our nucleation assay (Fig. 5b). In terms of fibril stability, all the functionally characterised mRIPK3 mutations decrease fibril stability, showing a positive correlation with ΔΔG (Fig. 5c, R=0.69, p=0.09). Lastly, transgenic mice expressing the mRIPK3 V448P variant showed resistance to necroptosis^33^. This variant both decreases amyloid nucleation (NS = – 0.78; Supp. Fig. 4g and h) and destabilizes the amyloid core (ΔΔG = 11.97 kcal/mol; Supp. Fig. 6h).

hRIPK1 mutations I539P, I541P and N545P decrease cell death when ectopically expressed in cells^4^ and all of them have low NS in our assay, decreasing amyloid nucleation in comparison to WT (Fig. 2a, Supp. Fig. 1e and Supp. Fig. 9a). In terms of stability, I539P and I541P destabilize the amyloid fold, while N545P can either stabilize or destabilize depending on the amyloid structure used to estimate the ΔΔG (Supp. Fig. 9b). Finally, the hRIPK1 A543D has been reported to have a WT-like function^34^. This variant presents a WT-like amyloid nucleation (Fig. 2a and Supp. Fig. 1e) and it is not-destabilizing in the 9HR6 structure (ΔΔG = 0.36 kcal/mol) but it is destabilizing in the 9HR9 and 8Z93 structures (ΔΔG = 2.17 and 9.18 kcal/mol, respectively).

To systematically assess the effect of novel RHIM variants in necroptosis, we set up a necroptosis assay in HeLa cells (Fig. 5d). These cells do not express hRIPK3^35^ but undergo necroptosis in response to appropriate stimuli upon exogenous expression of hRIPK3 (Supp. Fig. 10a and c). We generated stable HeLa cells expressing hRIPK3 or its variants under an inducible promoter (Fig. 5d) and measured their ability to sustain functional necroptosis by measuring cell-viability after TNF triggering (Fig. 5d). As a control, we evaluated the effect of the kinase dead mutant D142N and the RHIM dead mutant VQVG>AAAA (4A), finding that both abrogate necroptosis leading to improved cell viability compared to WT (Supp. Fig. 10a). By correlating the functional impact of the RHIM mutations (Supp. Fig. 10b) to NS and ΔΔG estimates (Fig. 5e-f) we found that novel mutations that exhibit WT-like amyloid nucleation also result in WT-like function (Fig. 5e). On the other hand, mutations that either decrease or increase amyloid nucleation tend to impair necroptosis (increase cell viability). This is evident for variants that have a big effect on amyloid nucleation as V458E (decreased nucleation) and G461Y (increased nucleation). Variants with milder effects on amyloid nucleation present only a partial loss of function (I452S and G457V). Lastly, D462A increases amyloid nucleation and maintains WT-like function. All these mutants also affect fibril stability, but the degree of destabilization varies based on the structure used to estimate the ΔΔG: I452S, G461Y and M468A are always destabilizing; D462A can either stabilize (Fig. 5f; PDB 7DAC) or destabilize (Supp. Fig. 8k-l; PDBs 7DA4 and 8Z94) the fibril structure; V458E is destabilizing in two of them (Fig. 5f and Supp. Fig. 8k; PDBs 7DAC and 7DA4) but is WT-like in one structure (Supp. Fig. 8l; PDB 8Z94) and L449A destabilizes only one of the fibril structures (PBD: 7DAC, Fig. 5f) but not the remaining two (PDBs 7DA4 and 8Z94, Supp. Fig. 8k-l). Evaluation of the hRIPK3 variants expression in transduced HeLa cells revealed that the effects on cell viability were not confounded by differential protein expression (Supp. Fig. 10c).

Taken together, these results suggest that disruption of amyloid nucleation, either by increasing or decreasing nucleation, as well as decreasing fibril stability, impairs necroptosis, suggesting that a “sweet-spot” of amyloid formation is needed for necroptosis.

### RIPK3 loss-of-function mutations do not form functional aggregates

To explore the subcellular distribution of the different hRIPK3 variants we performed immunostaining against RIPK3 after triggering necroptosis. Cells expressing the WT protein show dense fluorescent puncta corresponding to the fibrillar aggregates forming the necrosome (Fig. 5g), with a similar pattern as previously reported in the literature^22,25^. HeLa cells expressing hRIPK3 L449A or M468A mutations, both having a WT-like function (Fig. 5e-f and Supp. Fig. 10b), also form fluorescent puncta similar to those observed for cells expressing the WT protein. Puncta are also present in cells expressing hRIPK3 I452S, D462A, and G457V, but are smaller and occur less frequently (Fig. 5g). hRIPK3 V458E and G461Y, mutations that strongly perturb amyloid formation, decreasing and increasing it respectively, both lead to an almost complete loss of function in the viability assay (Fig. 5e and f and Supp. Fig. 10b). The signal for V458E is diffuse, while G461Y exhibits a predominantly diffuse signal with some cells featuring fluorescent puncta that may correspond to alternative non-functional aggregates (Fig. 5g). These data suggest that mutations strongly affecting amyloid formation in either direction also disrupt function.

### A “sweet-spot” of amyloid formation is required for signaling leading to necroptosis

We next asked whether variants affecting nucleation by the hRIPK1 and hRIPK3 RHIM domains are present in the human population. For this, we obtained allele frequencies for variants reported in GnomAD and in the Regeneron Genetics Center (RGC) database^36,37^ and correlated them with our NS and the estimated ΔΔGs. For both proteins, variants that severely affect nucleation are extremely rare in the human population (Fig. 6a and Supp. Fig. 8e-j and Supp. Fig. 9c-f). For hRIPK3 we also observe that variants with a WT-like NS are more frequent in the human population (Fig. 6a). In a similar manner, many of the variants that affect fibril stability are extremely rare in the human population, while those that are more frequent are WT-like or even increase fibril stability (Fig. 6b).

RIPK amyloids are functional, and their sequences contain evidence of evolutionary pressure (Fig. 1a and Supp. Fig. 1c). Our data suggest evolutionary optimization of the RHIM domain at a “sweet-spot” of amyloid propensity required for signaling. To relate evolutionary pressure with amyloid formation and amyloid stability, we annotated the NS and fibril ΔΔG landscape indicating mutations with reported functional effects on necroptosis and those that are more frequent in the human population (Allele Frequency (AF)>1e-5) (Fig. 6c and Supp. Fig. 11). All variants that decrease function cause alterations of ΔΔG above 2 kcal/mol in two of the reported hRIPK3 amyloid structures (PDBs 7DAC and 7DA4; Fig. 6c and Supp. Fig. 11a), suggesting they are incompatible with the amyloid fold required for function. These variants span across the entire range of nucleation scores. On the other hand, variants that are functional or are more frequent in the human population not only localize below the 2 kcal/mol threshold but also maintain a WT-like behaviour in terms of amyloid nucleation. When doing the same analysis but using the ΔΔGs estimated with the remaining structure (PDB 8Z94), we find that variants that decrease function also decrease amyloid nucleation or, as previously mentioned, have a ΔΔG above 2 kcal/mol (Supp. Fig. 11b). The only exception to this classification is M468D, which has a WT-like function but slightly increases amyloid nucleation and destabilizes the amyloid fold across the different structures.

## Discussion

Formation of RHIM-dependent RIPK1/RIPK3 amyloid filaments is required for full activation of necroptosis. Mutation of central residues in the RHIM domain alters amyloid aggregation and impacts necroptosis^4^. To date, the fibrillar architecture of RIPK1 and RIPK3 are the only human functional amyloids characterized at near-atomic resolution, positioning these sequences as a powerful model for mechanistic insight into how human cells can harness amyloid formation for signaling. However, only four mutations affecting the RHIMs have been characterized when it comes to the kinetics of amyloid formation (3 mRIP3 mutants, 1 hRIPK1)^19,21^ and only a few more for their impact on function, i.e. necroptotic cell death (4 hRIPK1, 13 hRIPK3 and 9 mRIPK3 mutants)^4,19,22,32,34^.

Here, we bridge the gap between structure and mechanism by performing a deep-mutational scan spanning ∼3,000 mutations across the RHIM domains and their flanking regions. By mapping how each variant alters amyloid formation we generate the first mutational landscape of a functional amyloid, revealing the determinants of RIPK assembly and defining how nucleation behaviour can encode functional outcomes in necroptosis.

Our mutational atlas revealed a conserved aliphatic tetrad in the first half of the RHIM domain that constitutes the first β-strand interface. Glycine residues emerged as key determinants of amyloid nucleation, with their substitution, as well as deletions in hRIPK3, typically enhancing aggregation. Notably, mRIPK3 carries a glutamic acid in place of a glycine at the position preceding the second β-strand, yet DMS shows its mutation mirrors the behaviour of equivalent glycines in other RHIMs. Interestingly, the ssNMR structure of mRIPK3 (PDB 6JPD) suggests that the local packing in this region is sufficiently open to accommodate this glutamic acid^19^.

Across the RIPKs studied here, mutations within the RHIM domain show similar behavior, as reflected by correlated NS effects, consistent with a conserved amyloid core. All reported RHIM homo-amyloid structures adopt a similar S-shaped fold formed by two β-sheet interfaces, and we find that mutations at residues contributing to these interfaces destabilize the fibrillar architecture.

In contrast, sequence-specific differences emerge in how individual β-sheet interfaces contribute to nucleation. In RIPK1, mutations in any β-strand tend to reduce amyloid nucleation. In RIPK3, however, only mutations in the first and second β-strands decrease nucleation, whereas mutations in the third β-strand have little effect or can even enhance nucleation. Flanking regions further diverge between sequences: the impact of mutations is highly sequence-dependent when it comes to these highly flexible regions, consistent with a role as entropic bristles that modulate solubility and actually prevent excessive aggregation thanks to gatekeeper residues.

Although the relevance of RHIM domain for RIPKs activity and necroptosis is known^4^, the molecular interplay between amyloid aggregation and kinase activity is still poorly understood. By relating the effect of RHIM mutations on necroptosis to their impact on amyloid nucleation (NS) and fibril stability (ΔΔG) we find that mutations destabilizing the fibril structure reduce necroptosis, as do mutations that either decrease or increase nucleation. Consistently, RIPKs mutations that affect either amyloid nucleation or fibril stability are extremely rare in the human population, whereas variants that preserve the nucleation propensity and do not reduce fibril stability are instead more common.

The data presented here suggest that necroptosis requires a narrow “sweet-spot” of amyloid formation. This sweet spot consists of an appropriate nucleation step that yields a stable amyloid fibril (Fig. 6b). Mutations that reduce nucleation may fail to form the necrosome or do so too slowly to support proper pathway signaling. Many of these mutations also destabilize the S-shaped fibril fold.

On the other hand, some mutations increase nucleation. When enhanced nucleation yields a stable S-shaped fibril, the necrosome forms and necroptosis proceeds. However, mutations that increase nucleation while destabilizing the S-shaped fold are unlikely to support necroptosis, likely because they promote alternative, non-productive amyloid conformations. Consistent with this and in contrast to pathological amyloids, RIPK functional amyloids are highly monomorphic^18–22^. Mutations such as hRIPK3 L466E and mRIPK3 L456D, which feature enhanced amyloid formation in our datasets, have indeed been shown to form distinct aggregate types in HeLa cells and *in vitro*, respectively^19,22^. Indels that increase nucleation in our data set may act similarly by altering backbone geometry and enabling alternative non-functional folds. Beyond aggregate fold, other factors may contribute to the functional outcome of RIPK variants. The heat shock protein family A member 8 (HSPA8) has been recently shown to disassemble RHIM-amyloids and inhibit necroptosis^34^ and could, for example, exhibit differential activity on different RIPK variants.

Necroptosis is a central cellular mechanism in host–pathogen interactions and in multiple human diseases^1^. Together, our findings suggest that functional necroptotic signaling relies on amyloid nucleation and fold identity operating within a narrow window, likely complemented by other layers of RIPK regulation. This sequence-resolved mechanistic understanding of the relationship between amyloid formation and function in RIPKs can help guide the development of therapeutic strategies that aim at modulating cell death. Beyond this, this work not only provides a framework for therapeutic modulation of necroptosis but also for engineering synthetic amyloids with tailored activities.

## Acknowledgements

Work in the lab of B.B. is supported by the la Caixa Research Foundation project ‘DeepAmyloids’ (LCF/PR/HR21/52410004), by the Spanish Ministry of Science, Innovation and Universities (PID2021-127761OB-I00, PID2024-157544OB-I00 and RYC2020-028861-I, funded by MCIN/AEI/10.13039/501100011033, “ERDF A way of making Europe” and “ESF Investing in your future”) and by the European Union (ERC Consolidator, Glam-MAP, 101125484). Views and opinions expressed are however those of the author(s) only and do not necessarily reflect those of the European Union or the European Research Council. Neither the European Union nor the granting authority can be held responsible for them. IBEC is a member of the CERCA Program/Generalitat de Catalunya. We thank the Chernoff lab for providing strains, James Murphy’s lab for sending the pTRE3G-FLAG-hRIPK3 vector, Fran Supek’s lab for pPAX2 and pVSVG plasmids. At IBEC, we thank Elena Martínez Fraiz’s lab for HT-29 cells and Irene Marco Rius’s lab for HeLa cells. Finally, we thank the CRG Genomics core technology for all the support with sequencing.

## Data availability

Raw sequencing reads are deposited in the European Nucleotide Archive (ENA) as part of the study PRJEB107723. The processed reads and nucleation scores are available as Dataset S3.

## Code availability

All scripts used for downstream analysis and to reproduce all figures are in https://github.com/BEBlab/DMS-RIPKs.

## Methods

### Library design and construction

Each designed library (hRIPK1, hRIPK3, mRIPK1, mRIPK3) contains a total of 1088 unique hRIPK1 nucleotide variants (680 single amino acid variants encoded by either 1, 2, and 3 nucleotide changes). The hRIPK3 substitutions and indels library also contains 1306 nucleotide variants (646 single amino acid variants encoded by 1, 2, and 3 nucleotide changes, 628 single amino acid insertions and 32 single amino acid deletions). The hRIPK1, hRIPK3, mRIPK1, mRIPK3 libraries were synthesized by Integrated DNA Technologies (IDT) as oligo pools covering the hRIPK1, hRIPK3, mRIPK1 and/or mRIPK3 102 nt sequence, flanked by 25 nt upstream and 21 nt downstream constant regions for cloning. The hRIPK3 substitutions and indels libraries were synthesized by Twist bioscience covering the hRIPK3 102 nt sequence, flanked by 24 nt upstream and 21 nt downstream constant regions for cloning. The hRIPK3 substitutions present in both libraries (IDT and Twist) were used for normalizing indels data (see section “Nucleation Scores and errors estimates”). For the IDT libraries, 2 ul of 10uM library pool were extended into a double stranded DNA by a single cycle PCR (Q5 high-fidelity DNA polymerase, NEB) with primers annealing to the 3’ constant region (primers MM_02, Dataset S1). For the Twist library, 20 ng of the library pool were amplified by a PCR of 12 cycles (Q5 high-fidelity DNA polymerase, NEB) with primers annealing to the constant regions (primers MM_01 and MM_02, Dataset S1). The products were purified from a 2% agarose gel (QIAquick Gel Extraction Kit, Qiagen). In parallel, the pCUP1-Sup35N plasmid was linearized by PCR (Q5 high-fidelity DNA polymerase, NEB; primers MM_03 and MM_04, Dataset S1). The product was purified from a 1% agarose gel (QIAquick Gel Extraction Kit, Qiagen). The libraries were then ligated into 200 ng of the linearized plasmid in a 1:10 (vector:insert) ratio by a Gibson approach with 3 h of incubation at 50°C followed by dialysis for 3 h on a membrane filter (MF-Millipore 0.025 μm membrane, Merck) and vacuum concentration. The product was transformed into 10-beta Electrocompetent *E. coli* (NEB), by electroporation at 2.0 kV, 200 Ω, 25 μF (BioRad GenePulser machine). Cells were recovered in SOC medium for 30 min and grown overnight in 50 ml of LB ampicillin medium. A small number of cells were also plated in LB ampicillin plates to assess transformation efficiency. A total of >100,000 transformants were estimated, meaning that each variant in each library is represented >100 times. 5 ml of overnight cultures were harvested to purify the RIPKs libraries with a mini prep (QIAprep Miniprep Kit, Qiagen).

### Yeast transformation

*Saccharomyces cerevisiae* [psi-pin-] (MATα ade1–14 his3 leu2-3,112 lys2 trp1 ura3–52) provided by the Chernoff lab was used in all experiments in this study. Yeast cells were transformed with the RIPKs libraries in three biological replicates. Per replicate, an individual colony was grown overnight in 3 ml YPDA medium at 30 °C and 4 g. Cells were diluted in 40 ml to OD600 = 0.3 and grown for 4–5 h. When cells reached the exponential phase (OD∼0.8–0.9) cells were harvested at 3000 × g for 5 min, washed with 50 ml milliQ, centrifuged at 3000 × g for 5 min and washed with 25 ml SORB buffer (100 mM LiOAc, 10 mM Tris pH 8.0, 1 mM EDTA, 1 M sorbitol). Cells were resuspended in 1.4 ml of SORB and incubated 30 min on an orbital shaker. After incubation, 800 ng of library and 30 ul of ssDNA (UltraPure, Thermo Scientific) were added and incubated for 5 min at room temperature and then 10 min on an orbital shaker. 6 ml of YTB-PEG (100 mM LiOAc, 10 mM Tris pH 8.0, 1 mM EDTA, 40% PEG 3350) and 580 ul of DMSO were added to the cells. Heat-shock was performed at 42°C for 20 min in a liquid bath with intermittent shaking. Cells were harvested and incubated in 50 ml of recovery medium (YPDA medium + Sorbitol 0.5 M) for 1 h at 30°C. Cells were harvested and grown in 50 ml plasmid selection medium (-URA, 2% glucose) for 50 h at 30 °C. A small amount of cells were also plated in plasmid selection solid medium to assess transformation efficiency. 50,000-175,000 transformants were estimated for each biological replicate, meaning that each variant in the library is represented at least 50 times. After 50 h, cells were diluted in 50 ml plasmid selection medium to OD= 0.05 and grown exponentially for 15 h. Finally, the culture was harvested and stored at −80 °C in 25 % glycerol.

### Selection experiments

*In vivo* selection assays were performed in three independent biological replicates for each RIPK library. For each replicate, cells were thawed from −80 °C in 50 ml plasmid selection medium at OD = 0.05 and grown until exponential for 15 h. At this stage, cells were harvested and resuspended in 50 ml protein induction medium (-URA, 2% glucose, 100 μM Cu2SO4) at OD = 0.1. After 24 h the 40 ml input pellets were collected, and cells were plated on –ADE-URA selection medium in 145-cm^2^ plates (Nunc, Thermo Scientific). Plates were incubated at 30 °C for 7 days. Finally, colonies were scraped off the plates with PBS 1x and harvested by centrifugation to collect the output pellets. Both input and output pellets were stored at −20 °C before DNA extraction. Three input and three output samples were processed for sequencing.

### DNA extraction and sequencing library preparation

Input and output pellets were resuspended in 0.5ml extraction buffer (2% Triton-X, 1% SDS, 100 mM NaCl, 10 mM Tris-HCl pH 8, 1 mM EDTA pH 8). They were then frozen for 10 min in an ethanol-dry ice bath and heated for 10 min at 62 °C. This cycle was repeated twice. 0.5ml of phenol:chloroform:isoamyl (25:24:1 mixture, Thermo Scientific) was added together with glass beads (Sigma). Samples were vortexed for 10 min and centrifuged for 30 min at 2000 × g. The aqueous phase was then transferred to a new tube, and mixed again with phenol:chloroform:isoamyl, vortexed and centrifuged for 45 min at 2000 × g. Next, the aqueous phase was transferred to another tube with 1:10 V 3 M NaOAc and 2.2 V cold ethanol 96% for DNA precipitation. After 30 min at −20 °C, samples were centrifuged and pellets were dried overnight. The following day, pellets were resuspended in 0.3 ml TE 1X buffer and treated with 10 μl RNAse A (Thermo Scientific) for 30 min at 37 °C. DNA was finally purified using 10 μl of silica beads (QIAEX II Gel Extraction Kit, Qiagen) and eluted in 30 μl elution buffer. Plasmid concentrations were measured by quantitative PCR with SYBR green (A25742, Applied biosystems) and primers annealing to the origin of replication site of the pCUP1-Sup35N plasmid at 58 °C for 40 cycles (primers MM_05-06, Dataset S1). The library for high-throughput sequencing was prepared in a two-step PCR (Q5 high-fidelity DNA polymerase, NEB). In PCR1, 50 million of molecules (RIPKs libraries) were amplified for 15 cycles with frame-shifted primers with homology to Illumina sequencing primers (primers MM_07–20, Dataset S1). The products were treated with ExoSAP treatment (Affymetrix) and purified by column purification (MinElute PCR Purification Kit, Qiagen). They were then amplified for 12 cycles in PCR2 with Illumina-indexed primers (primers MM_21–99, Dataset S1). For each of the RIPKs libraries, the six samples (one input-output pair per biological replicate) were pooled together equimolarly and the final product was purified from a 2% agarose gel with 20 μl silica beads (QIAEX II Gel Extraction Kit, Qiagen). The libraries were sent for 125 bp paired-end sequencing in an Illumina NextSeq500 sequencer at the CRG Genomics core facility. In total, >10 million paired-end reads were obtained for each of RIPKs libraries, representing >1000x read per variant coverage.

### Individual variant testing

Human and mouse RIPK1 and RIPK3 sequences (primers MM_100-103, Dataset S1) were bought as ultramers from IDT and 2 ul of a 10uM solution of each ultramer were used as a template to extend the sequence into a double stranded DNA by a single cycle PCR (Q5 high-fidelity DNA polymerase, NEB) with primers annealing to the 3’ constant region (primers MM_02, Dataset S1). Human RIPK2 cDNA was obtained from the human orfeome library (Protein Techonologies Unit, CRG) and amplified with primers (MM_104 and MM_105, Dataset S1) with homologous regions to the pCUP1-Sup35N vector. The products were cloned into the pCUP1-Sup35N vector with the same protocol used as for the libraries. Selected variants were obtained via PCR mutagenesis (primers MM_106-132). Plasmids were transformed into *E. coli* and then yeast, with mutations verified by Sanger sequencing. Yeast expressing each variant were grown in induction media (-URA 2% glucose 100 μM Cu2SO4), then plated on control (-URA) and selective (-URA-ADE) plates for colony counting.

### Cell lines and cell culture

HT29, HEK293T and HeLa cells were obtained from collaborators at IBEC (Elena Martínez Fraiz for HT-29 cells, Elisabet Engel for HEK293T and Irene Marco Rius for HeLa cells). All cell lines were maintained at 37 °C and 5% CO2 in Dulbecco’s modified Eagle’s medium (DMEM-HG; Gibco, 11668027) supplemented with 10% fetal bovine serum (FBS; Gibco, A5256801) and 100 units/mL of penicillin and 100μg/ml streptomycin (P/S; Gibco, 15140122). This media was named as D10 medium. Cells were routinely tested to be free of mycoplasma contamination.

### Constructs and transfection

For lentivirus production, the WT and mutated hRIPK3 cDNAs were cloned into the modified lentiviral vector pTRE3G. HEK293T cells were seeded on multiwell-6 cm dishes and cultured to 70% confluence in D10 medium. The cells were then transfected with the lentiviral vectors and virus packing plasmids (psPAX2 and pVSVG) by Lipofectamine™ 2000 (11668027, Invitrogen) according to the manufacturer protocol. The virus containing medium was harvested 48 and 72 hs later.

HeLa cells grown until 70% confluency in D10 medium were transduced with the virus containing media and 8 μg/ml polybrene (TR-1003, Sigma Aldrich). The infection medium was changed with fresh D10 medium 24 hs later. Selection of the transduced cells with 1 ug/ml of Puromycin (SC108071-SC, Santa Cruz) started 48 hs after transduction. The selected cells were maintained in Puromycin until reaching confluency.

### Cell survival assay

Transduced Hela cells with lentiviral particles carrying the wild-type or mutant hRIPK3 cDNA were cultured to 90% confluence, before trypsin digestion and seeding in black 96-well plates (137101, Thermo Scientific). 5000-7000 cells were seeded in each well, including triplicate wells (technical replicates). After 24 hs, hRIPK3 expression was induced with 20 ng/ml of Doxycycline (D9891, Sigma Aldrich). After 24 hs, necroptosis was triggered by adding TNF-α at a final concentration of 100 ng/ml (T, H8916, Sigma Aldrich), Brinipant at 50 nM (Smac mimetic (S), HY-16591, MedChemExpress) and Emricasan at 10 μM (Pancaspase inhibitor (I), HY-10396, MedChemExpress) to the cell culture wells. After 24 hs, cell survival was determined by measuring cellular ATP level with the Cell Titer-Glo Luminescent Cell Viability Assay kit (G7571, Promega) according to the manufacturer’s instructions. Luminescence was measured with a Spark Multimode Plate Reader from TECAN. Each cell viability measurement was obtained in three independent replicates (from independently transduced HeLa cells).

### Immunofluorescence

Transduced Hela cells were seeded in glass coverslips. 24 hs later hRIPK3 expression was induced with 2 µg/ml of Doxycycline (D9891, Sigma Aldrich). After 24 hs, necroptosis was triggered as previously described. After 4 hs of incubation, cells were fixed with paraformaldehyde 4% (15710, Electron Microscopy Sciences 15710) for 10-15 min at room temperature. After three washes with PBS, cells were blocked-permeabilized with PBS + 2% BSA (A4503, Sigma) + 0.1% TritonX-100 (T8787, Sigma) for 1 hr at room temperature. Cells were incubated with anti-RIP3 (sc-374639, Santa Cruz) 1:250 in PBS + 0.2% BSA + 0.01% TritonX-100 ON at 4°C. The day after, cells were washed three times with PBS and incubated with the secondary antibody anti-mouse 488 (A-11001, Invitrogen) for 1 hr at room temperature. Finally cells were incubated with PBS + Dapi for 5 min at room temperature and washed two times with PBS. The coverslips were mounted using Vectashield Plus Mounting Media (H-1900-2, Vectashield). Immunofluorescence images were taken with Leica AF7000.

### Western Blot

Transduced Hela cells were seeded in 12-well plates. 24 hs later hRIPK3 expression was induced with 2 µg/ml of Doxycycline (D9891, Sigma Aldrich). After 24 hs, RIPK3 expressing cells were harvested in 2x Loading Buffer (4% SDS, 20% Glycerol, 120mM Tris HCl pH 6.8, 10% beta-mercaptoethanol and Bromophenol blue) and boiled at 95°C for 5, and then resolved by 4–12% Tris-Glycine gel (MB46501, NZYtech). Proteins were transferred to a nitrocellulose membrane and blocked with 5% w/v skim milk powder in PBS. Membranes were probed with antibodies against RIPK3 1:500 (sc-374639, Santa Cruz) and GAPDH 1:2000 (ab125247, Abcam) and secondary antibodies against mouse couple to HRP 1:5000 (sc-516102, Santa Cruz), and revealed using ECL Prime (GERPN2232, Cytiva).

### Data processing

FastQ files from paired end sequencing of the libraries were processed using DiMSum (https://github.com/lehner-lab/DiMSum)^38^, an R pipeline for analyzing deep mutational scanning data. 5′ and 3′ constant regions were trimmed, allowing a maximum of 20% of mismatches relative to the reference sequence. Sequences with a Phred base quality score below 30 were discarded. Non-designed variants were also discarded for further analysis, as well as variants with input reads below the threshold suggested by DimSum in all of the replicates.

### Nucleation scores and error estimates

The DiMSum package (https://github.com/lehner-lab/DiMSum)^38^ was also used to calculate nucleation scores (NS) and their error estimates for each variant in each biological replicate as:

Nucleation score = ESi – ESwt

Where ESi = log(Fi OUTPUT) – log(Fi INPUT) for a specific variant and ESwt = log(Fwt OUTPUT) – log(Fwt INPUT) for each RIPK.

NSs for each variant were merged across biological replicates using error-weighted mean and centered to the WT NS. All NS and associated error estimates are available in Dataset S3.

All RIPKs libraries were centered using the corresponding synonymous NS (except for the hRIPK3 indels library that was centered on WT estimates). The hRIPK3 substitutions present in both libraries (IDT and Twist libraries) were used for normalizing indels data. After NSs centering, each variant was tested against the WT NS at FDR=0.1 and at FDR=0.01 and classified in three possible groups: WT-like, NS_inc and NS_dec. Sigma values were normalized to the interquartile range and WT-like variants with a normalized sigma value above a cut-off of 0.3 were considered as noisy variants and excluded from further analysis. NS was obtained for 673 unique hRIPK1 variants, 1305 unique hRIPK3 variants, 657 unique mRIPK1 variants and 677 mRIPK3 unique variants. 667 confident estimates (with low normalized sigma value) were obtained for the hRIPK1 library, 588 for the hRIPK3 library, 652 for the mRIPK1 library and 660 for the mRIPK3 library. All NS estimates are available in Dataset S3.

### Fibril stability analysis and 3D protein visualization

Following the example of a recent study on PI3K-SH3 amyloids^39^, where FoldX^31^ predictions of ΔΔG were shown to correlate well with *in vitro* measurements of fibrils stability for 15 PI3K-SH3 variants (26), FoldX was ran on a stacked single filament trimer fibril structures of RIPKs peptides. For PDB files containing more than three chains, the remaining chains were deleted with the aim of running FoldX on trimeric fibril structures only. For the 9HR9 structure that contains only a single chain, the trimer was obtained by replicating this chain and by imposing a structural alignment with the trimeric structure of 9HR6 using the MultiSeq tool of VMD^40^. According to the FoldX manual, before calculating the ΔΔG, a “Repair PDB” step was run on the trimeric structures. The ΔΔG estimation was obtained running the “Build Model” command on FoldX with “numberOfRuns”=5 and performing the same single amino acid substitution in all the chains. ΔΔG values above 2 kcal/mol were considered as highly destabilizing. 3D protein visualization was done with VMD and Chimera^39–41^.

For hRIPK1, the predicted effects of mutating residue Y531 differ strikingly among structures. In the structure 9HR6 most mutations are destabilizing, in 8Z93 most mutations are stabilizing or WT-like, while in 9HR9 most mutations are stabilizing. One possible explanation is that after repairing the PDB with FoldX, residue Y531 in one of the monomers in structures 8Z93 and 9HR9 presents a different side chain orientation compared to the other monomers. Despite this technical problem we decided to include these results.

### Variant Effect Predictors

The VEPs scores were retrieved from https://huggingface.co/spaces/ntranoslab/esm_variants^29^, https://alphamissense.hegelab.org/^42^ and https://www.lcqb.upmc.fr/GEMME/^30^.

### Reported data and correlations

The effect of different RHIM mutations in necroptotic cell death or aggregation kinetics was obtained from previous literature ^4,19,22,32,34^. We used published Source Data Files when available. In case the data was not published as an accessible table, we extracted the values from the plots using Automeris (https://automeris.io/). Gnomad and Regeneron Genetics Center (RGC) data (reference transcript: NM_006871.4) was retrieved from the webpages on 30th October 2025^36,37^. The Regeneron Genetics Center, and its collaborators (collectively, the “Collaborators”) bear no responsibility for the analyses or interpretations of the data presented here. Any opinions, insights, or conclusions presented herein are those of the authors and not of the Collaborators. This research has been conducted using the UK Biobank Resource under application number 26041.

### Statistics and reproducibility

Based on transformation efficiency, each variant in the designed libraries (n=1088 nucleotide variants per library) is expected to be represented at least 50x, a coverage which was maintained at each step of the selection experiments and in library preparation for sequencing. In terms of sequencing, reads that did not pass the QC filters using the DiMSum package were excluded (https://github.com/lehner-lab/DiMSum)^38^. The experiments were not randomized. The Investigators were not blinded to allocation during experiments and outcome assessment.

## Figures legends

**Supplementary Figure 1.**
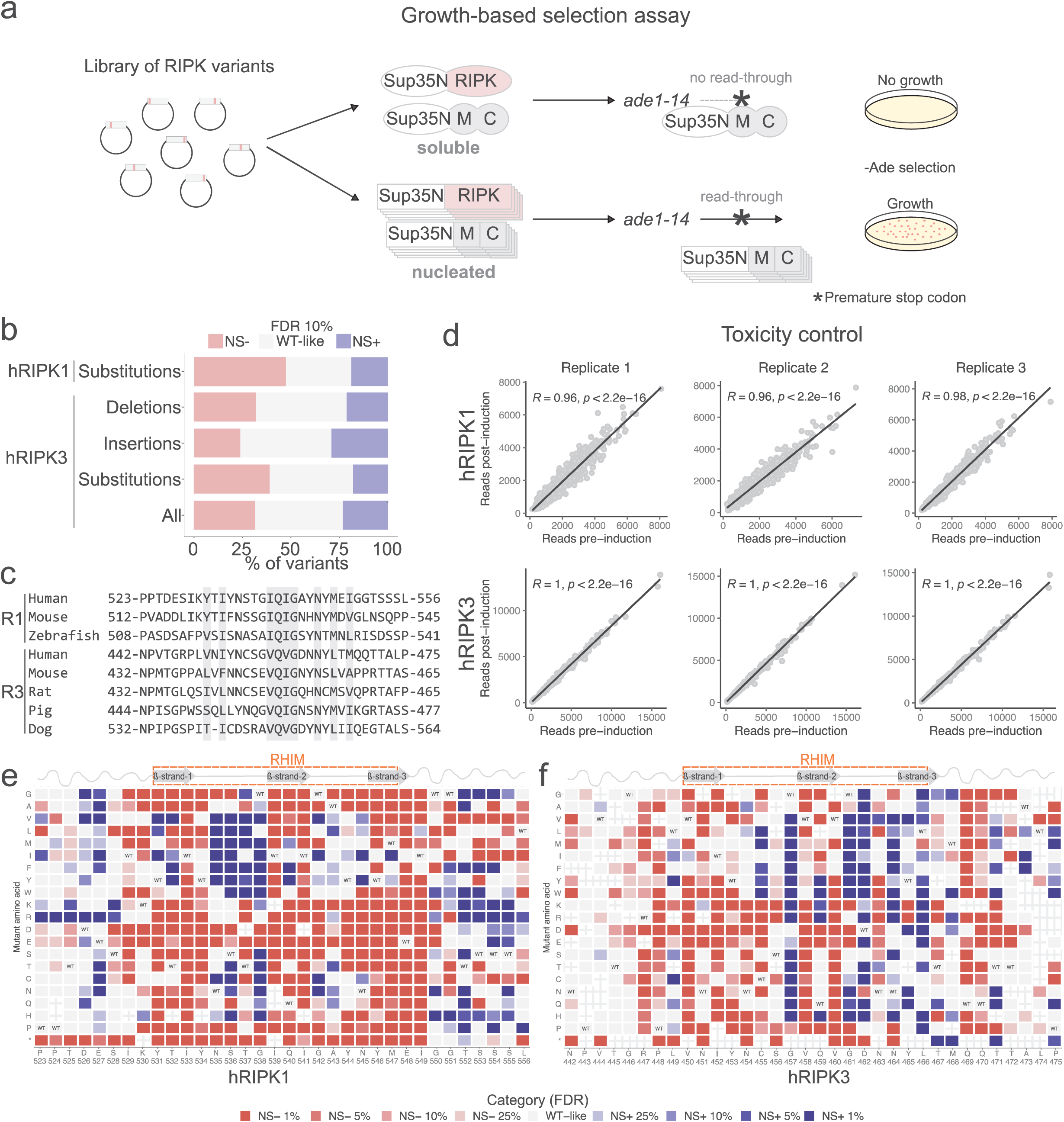
Deep mutagenesis of human hRIPK1 and hRIPK3. **a.** Percentage of variants increasing or decreasing nucleation at FDR = 0.1 and variants that are WT-like in the hRIPK1 and hRIPK3 libraries. **b.** RIPK1 and RIPK3 multiple sequence alignment performed by ClustalO^43^. Conserved residues are shaded. **c.** Toxicity control for hRIPK1 (top) and hRIPK3 (bottom) libraries. Deep sequencing is performed before and after 24h of protein expression. Pearson’s correlation coefficients and p-values are indicated. **d-e.** Heatmap of FDR categories for the effects of single amino acid substitutions of hRIPK1 (d) and hRIPK3 (e). The WT amino acid and position are indicated in the x-axis and the mutant amino acid is indicated in the y-axis. Synonymous mutations are indicated with “WT”.

**Supplementary Figure 2.**
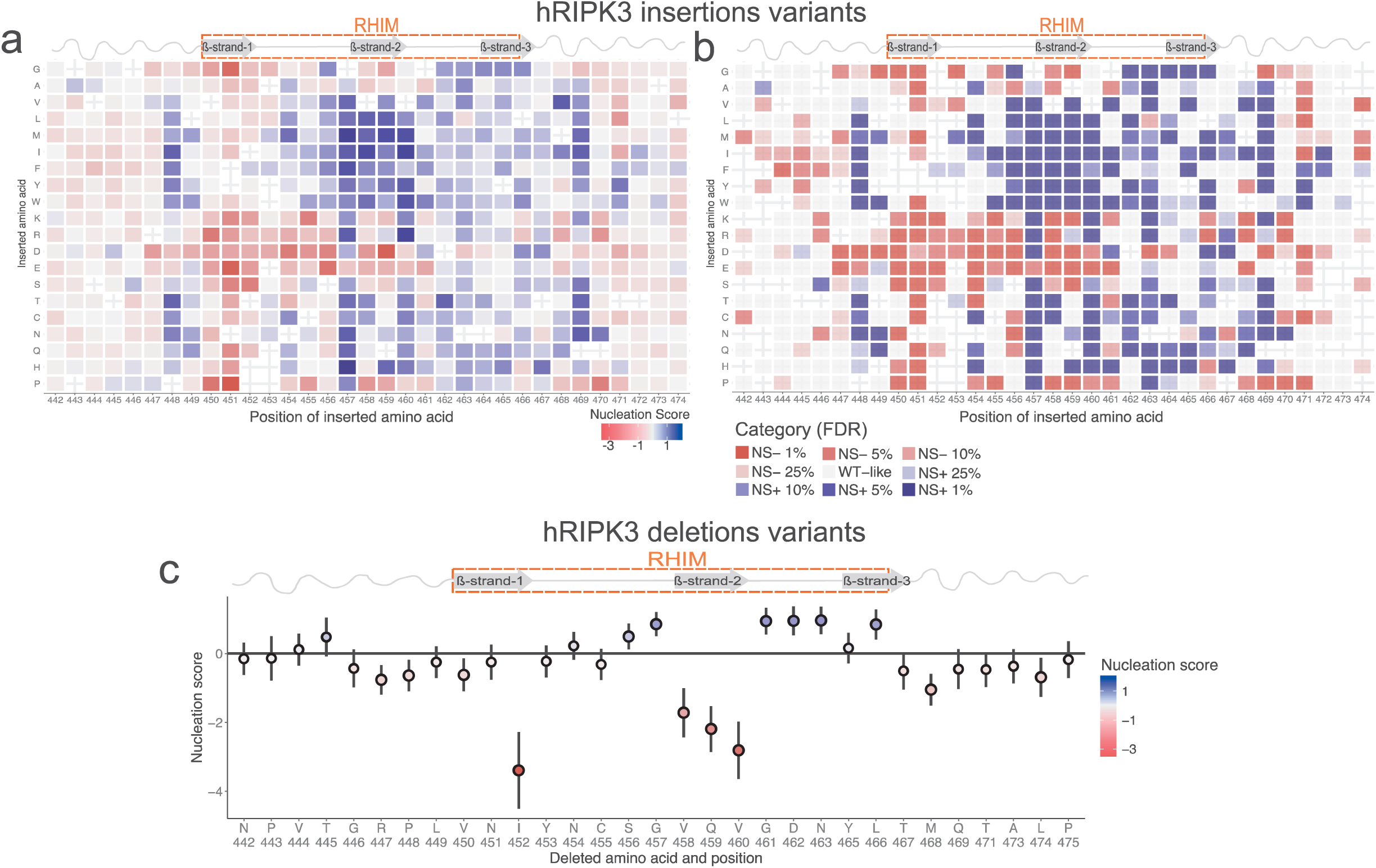
Deep indel mutagenesis of hRIPK3. **a.** Heatmap of nucleation scores for hRIPK3 single amino acid insertions. **b.** Heatmap of FDR categories for the effects of single amino acid insertions in hRIPK3 sequence. The WT amino acid and position are indicated in the x-axis and the mutant amino acid is indicated in the y-axis. Synonymous mutations are indicated with “WT”. **c.** Nucleation scores for RIPK3 single amino acid deletions. Vertical error bars indicate 95% CI of the mean. Circles with shaded external lines indicate significance at FDR=0.1.

**Supplementary Figure 3.**
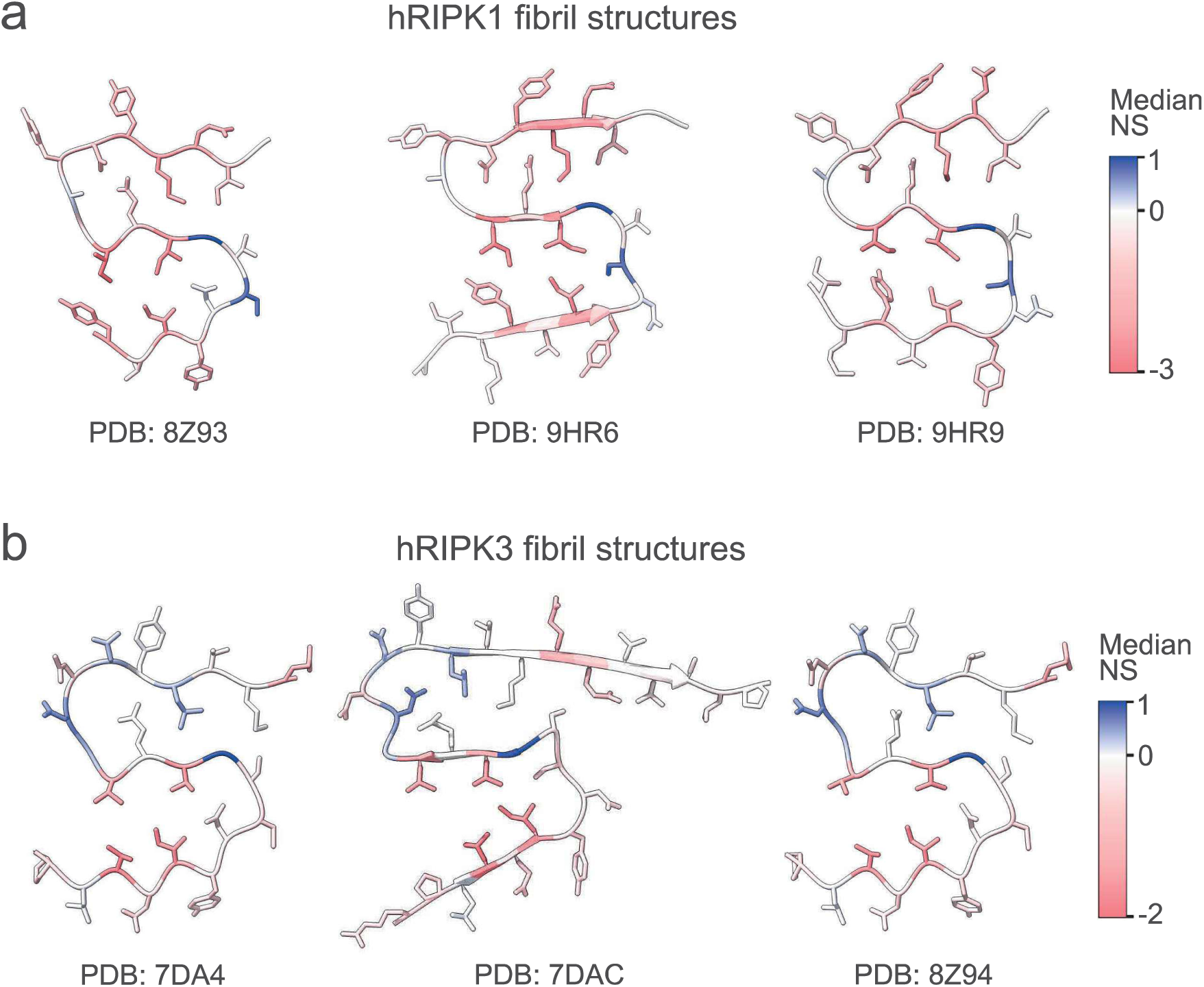
Monomeric cross-sections of hRIPK1 (a) and hRIPK3 (b) amyloid fibrils coloured by median nucleation score per position^18–22^.

**Supplementary Figure 4.**
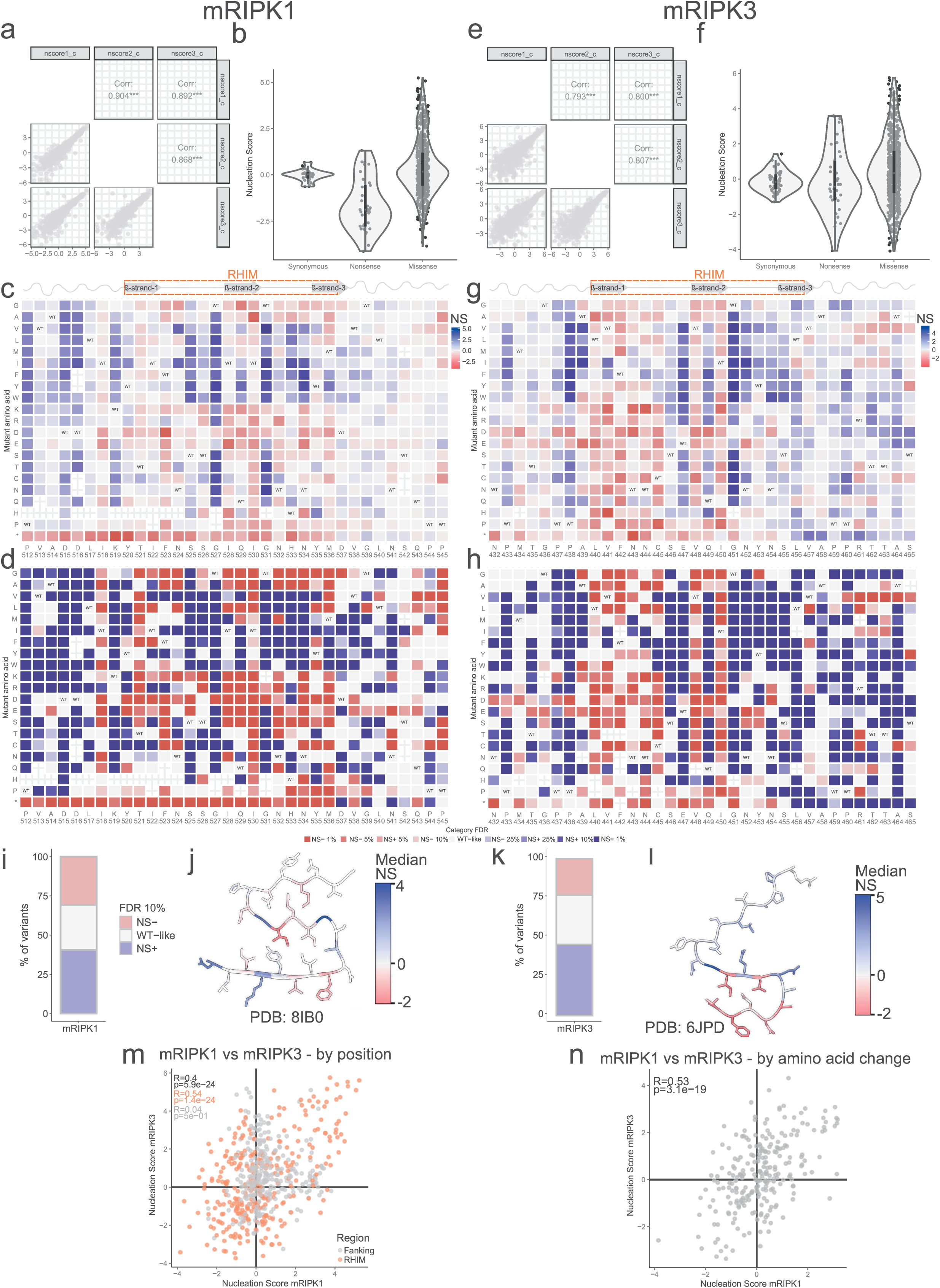
Deep mutagenesis of mouse RIPK1 and RIPK3. **a** and **e.** Correlation of nucleation scores across biological replicates for variants in the mRIPK1 library (a) and mRIPK3 library (e). Pearson’s correlation coefficients are indicated. **b** and **f.** Distribution of nucleation scores for each class of mutations in mRIPK1 library (b) and mRIPK3 library (f). **c** and **g.** Heatmap of nucleation scores for mRIPK1 (c) and mRIPK3 (g) single amino acid substitutions. The WT amino acid and position are indicated in the x-axis and the mutant amino acid is indicated in the y axis. Synonymous variants are indicated as “WT”. **d** and **h.** Heatmap of FDR categories for the effects of single amino acid substitutions of mRIPK1 (d) and mRIPK3 (h). The WT amino acid and position are indicated in the x-axis and the mutant amino acid is indicated in the y-axis. Synonymous mutations are indicated with “WT”. **i** and **k.** Percentage of variants increasing or decreasing nucleation at FDR = 0.1 and variants that are WT-like in the mRIPK1 (i) and mRIPK3 (k) libraries. **j** and **l**. Monomeric cross-section of mRIPK1 (j) and mRIPK3 (l) amyloid fibril coloured by median nucleation score per position. **m-n.** Correlation of mRIPK1 and mRIPK3 nucleation scores by position (m) or by amino acid change (n). Pearson’s correlation coefficients and p-values are indicated in black for the whole mutagenized sequence, in orange for the RHIM domain and in grey for the flanking regions.

**Supplementary Figure 5.**
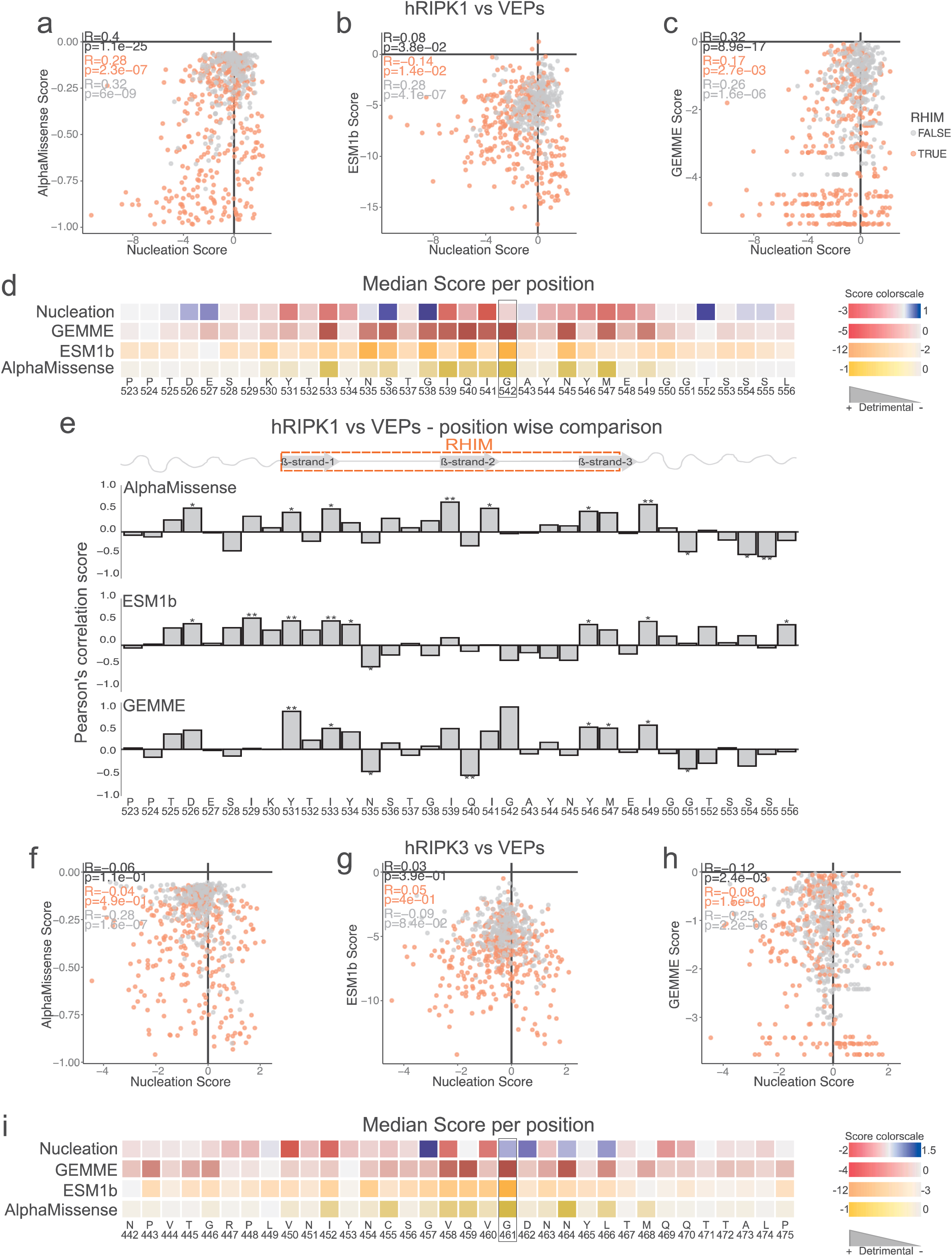
Comparison of hRIPK1 and hRIPK3 nucleation scores with variant effect predictors (VEPs) scores. **a-c and f-h.** Correlation of VEPs scores (Alphamissense a and f, ESM1b b and g, GEMME c and h)^28–30,42^ with hRIPK1 (a-c) and hRIPK3 (f-h) nucleation scores. Pearson’s correlation coefficients and p-values are indicated in black for the whole mutagenized sequence, in orange for the RHIM domain and in grey for the flanking regions. **d and i**. Comparison of median scores per position across nucleation score and VEPs scores for hRIPK1 (d) and hRIPK3 (i). Glycine residues with highest detrimental VEPs scores are highlighted with a black square. **e.** Position based correlation of hRIPK1 nucleation scores versus Alphamissense (top), ESM1b (middle) and GEMME (bottom) scores. Pearson’s correlation coefficient is indicated in the y-axis. p-value: *<0.05, **<0.01 and ***<0.001.

**Supplementary Figure 6.**
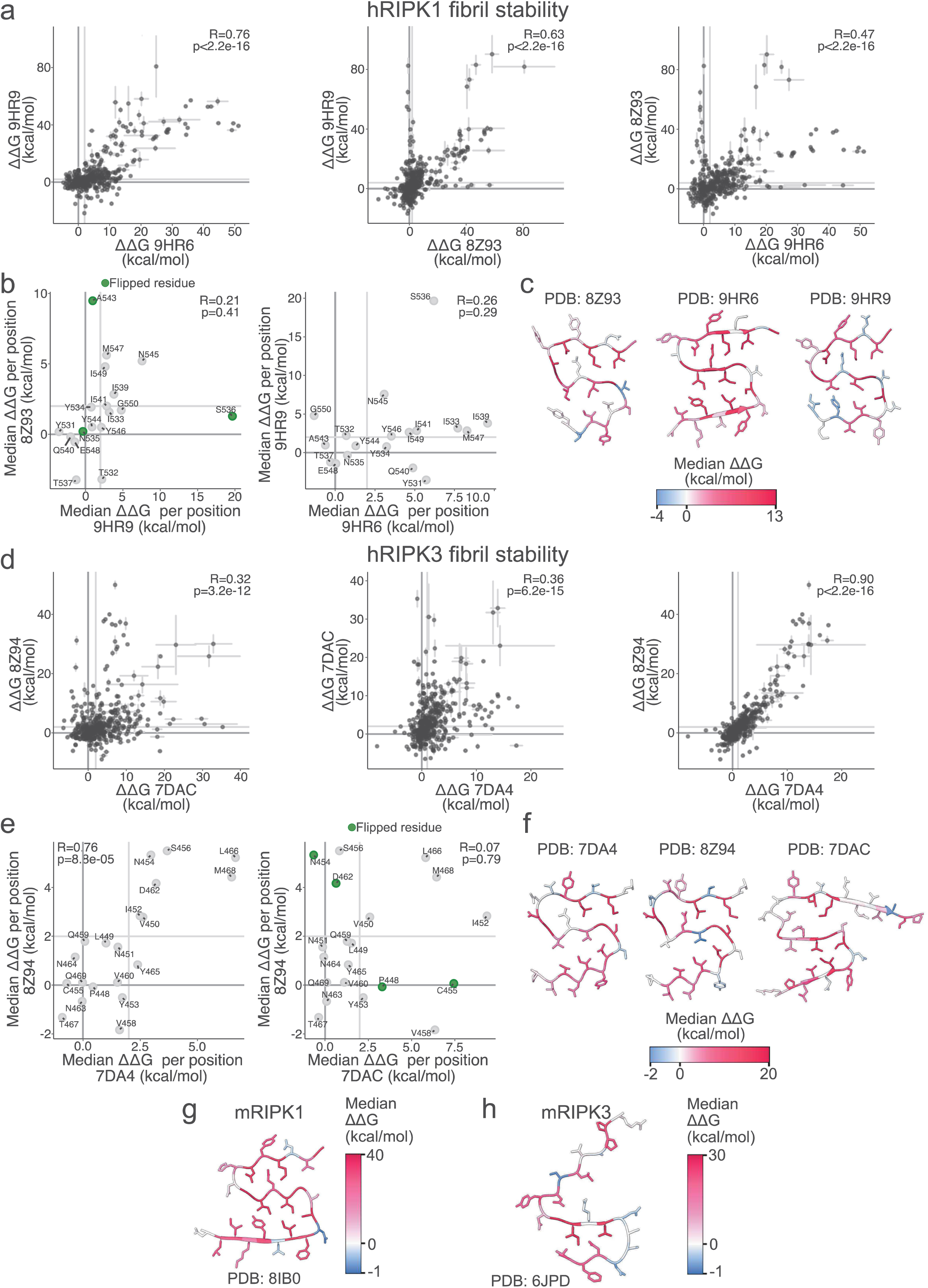
Effects of mutations on RIPKs fibrils stability. **a** and **d.** Correlation of ΔΔG estimates from FoldX between different hRIPK1 (a) and hRIPK3 (d) amyloid structures. **b** and **e.** Correlation of median ΔΔG estimates per position between different hRIPK1 (b) and hRIPK3 (d) structures. Pearson’s correlation coefficients and p-values are indicated. **c-h.** Monomeric cross-section of hRIPK1 (c), hRIPK3 (f), mRIPK1 (g) and mRIPK3 (h) amyloid fibrils coloured by median ΔΔG estimates per position.

**Supplementary Figure 7.**
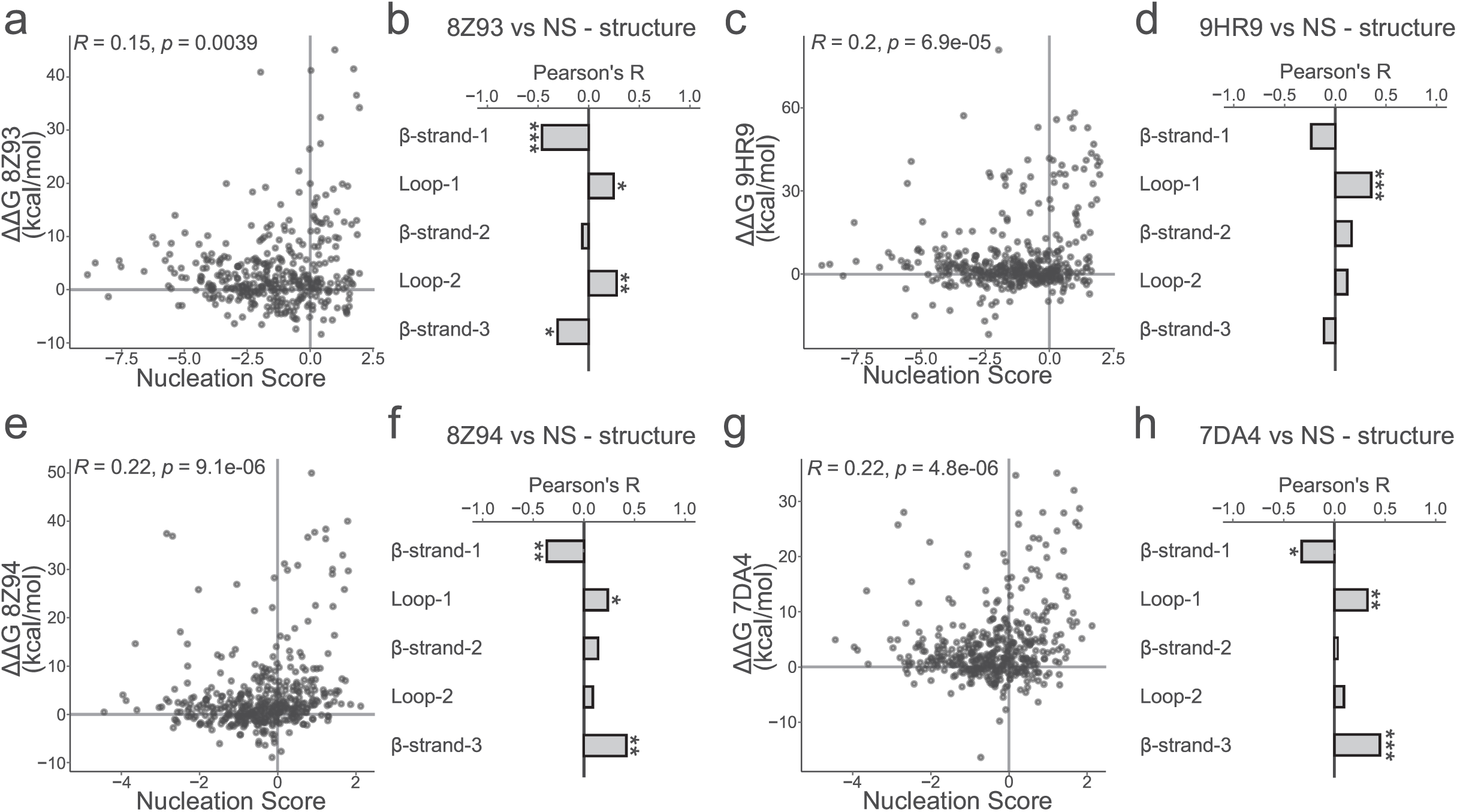
Correlation between nucleation scores and ΔΔG estimates. **a, c, e** and **g.** Correlation between hRIPK1 (a and c) and hRIPK3 (e and g) ΔΔG estimates and nucleation scores per residue change. **b, d, f** and **h.** Nucleation scores and ΔΔG correlations for hRIPK1 (b and d) and hRIPK3 (f and h) structural elements in the S-shaped fold. Pearson’s correlation coefficients and p-values are indicated. p-value: *<0.05, **<0.01 and ***<0.001.

**Supplementary Figure 8.**
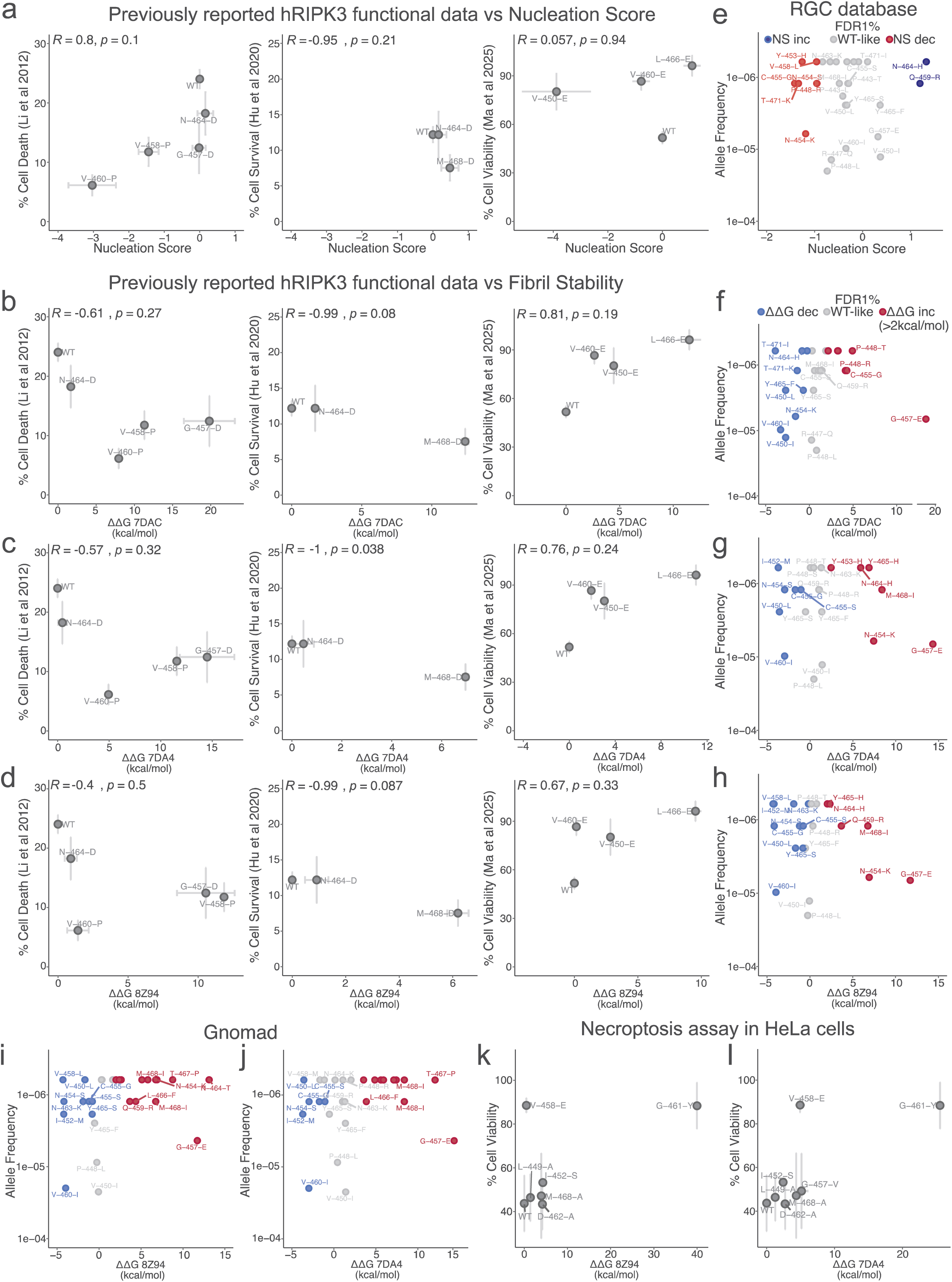
Variants altering hRIPK3 amyloid formation disrupt necroptosis. **a-d.** Correlation between previously reported cell death, cell survival or cell viability measurements and nucleation scores (a) or FoldX ΔΔGs (b-d) for hRIPK3 variants. Vertical and horizontal error bars indicate SD (n=3) and sigma errors of the experiments (in the case of the nucleation score) or SD from five FoldX runs (in the case of fibril stability estimates). **e-j.** hRIPK3 variants allele frequency from RGC^36^ (e-h) or Gnomad^37^ (i-j) versus nucleation scores (e) and FoldX ΔΔGs (f-j). **k-l.** Correlation between cell viability measurements from this study and ΔΔGs for hRIPK3 variants estimated using different amyloid structures (PDB: 7DAC, 7DA4 and 8Z94)^20,22^. Vertical and horizontal error bars indicate estimated SD (n=3) and sigma errors of the experiments (in the case of the nucleation score) or SD from five FoldX runs (in the case of fibril stability estimates).

**Supplementary Figure 9.**
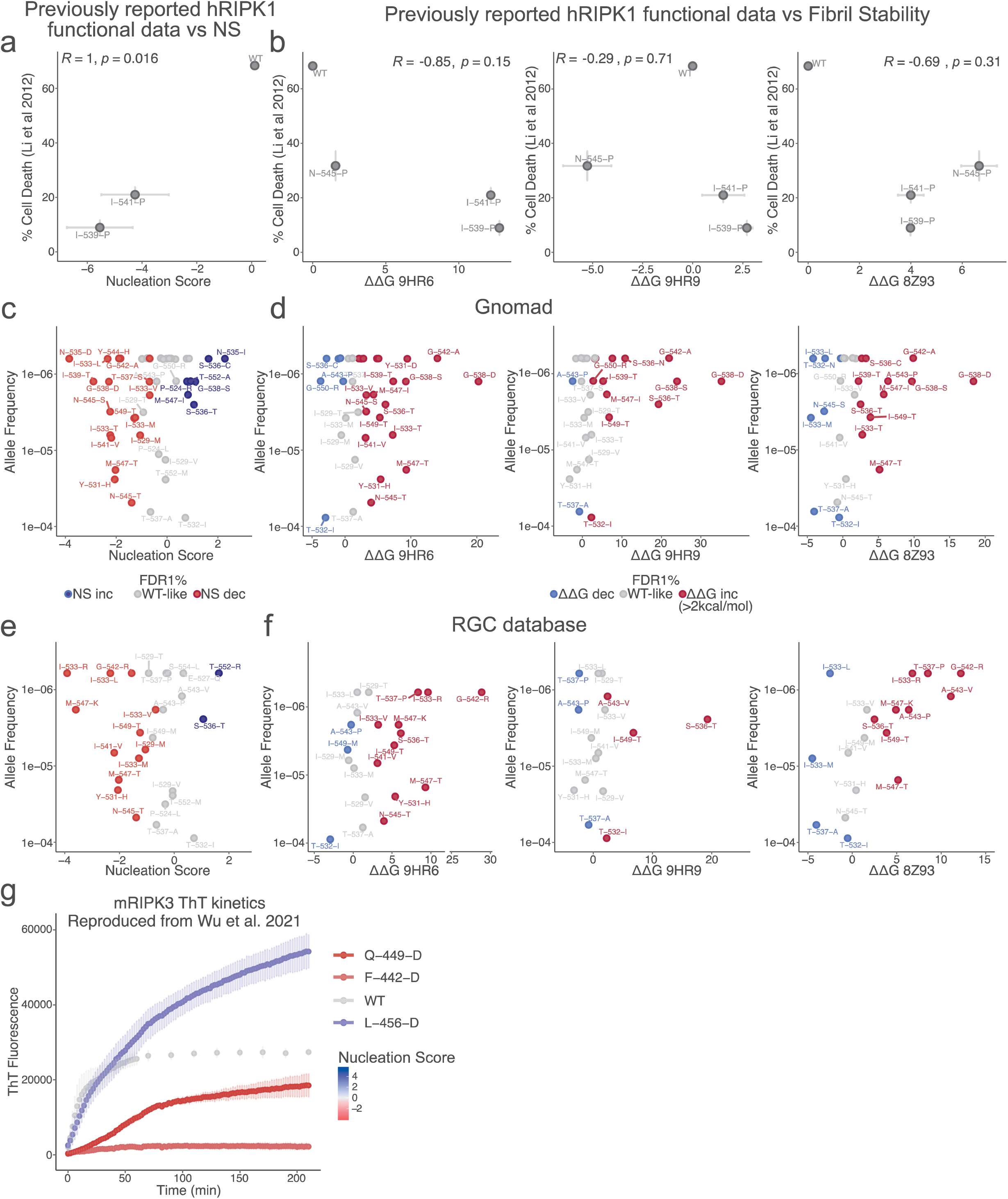
RIPKs amyloid formation and necroptosis. **a-b.** Correlation between cell death and nucleation scores (a) or FoldX ΔΔGs (b) for hRIPK1 variants. Vertical and horizontal error bars indicate estimated SD (n=3) and sigma errors of the experiments (in the case of the nucleation score) or SD from five FoldX runs (in the case of fibril stability estimates). **c-d.** hRIPK1 variants allele frequency from Gnomad versus nucleation scores (c) and ΔΔG (d). **e-f.** hRIPK1 variants allele frequency from RGC versus nucleation scores (c) and ΔΔG (d). **g.** Kinetics of amyloid formation of mRIPK3 and its variants reproduced from Wu et al. 2021^19^. The line of each trace is coloured based on nucleation scores measured in this study (blue: NS increased, grey: wt-like and red: NS decreased).

**Supplementary Figure 10.**
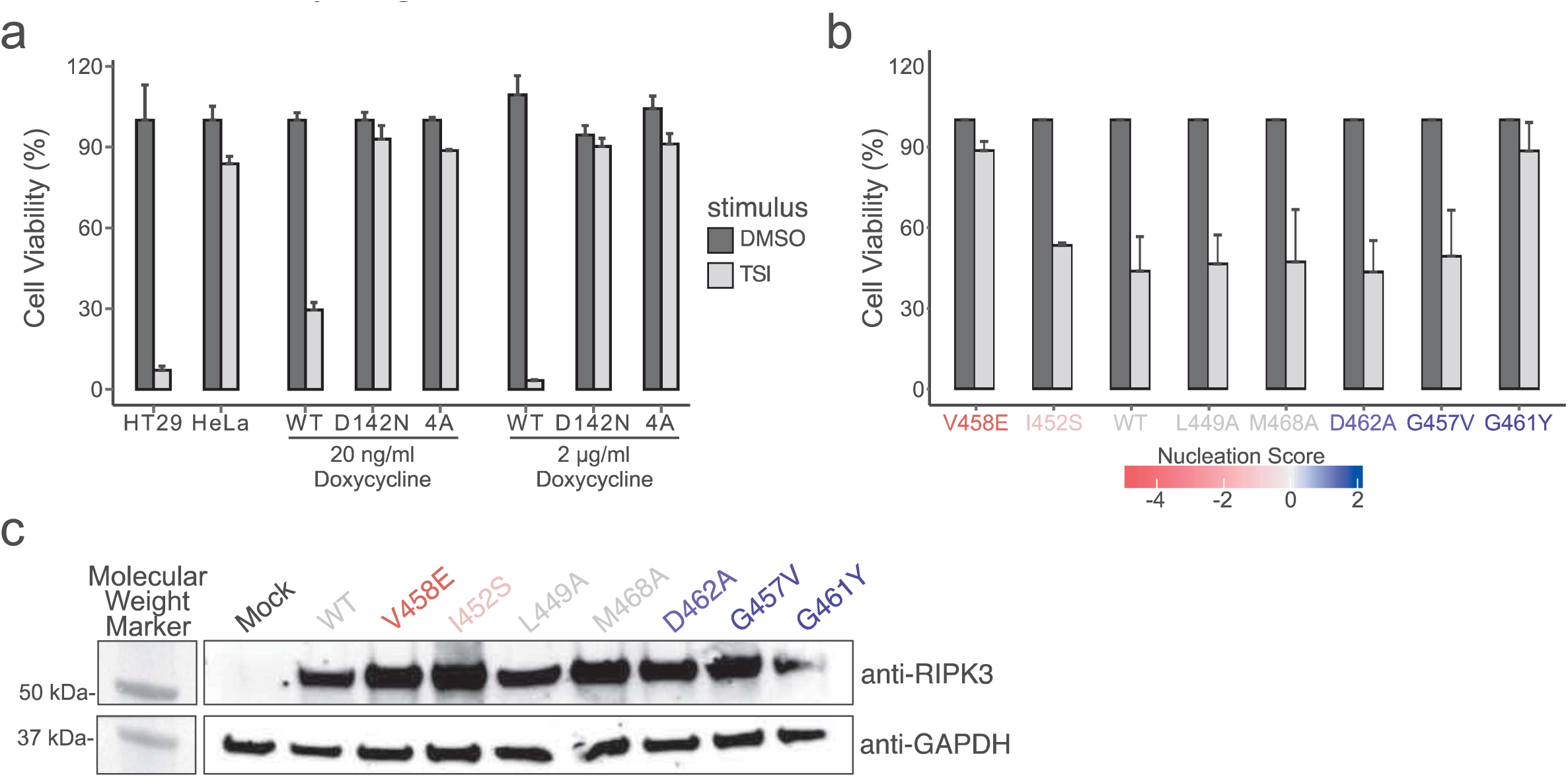
Effect of hRIPK3 mutations on cell viability. **a** and **b.** Cell viability (%) measured upon stimulation with TSI or with no stimulus (DMSO). The expression of hRIPK3 and variants was induced with two different concentrations of Doxycycline in HeLa transduced cell lines. The HT29 cell line was used as a positive control for the TSI stimulus. Variants known to disrupt necroptosis were used as controls (a) and the functional outcome of mutations in the RHIM domain prioritised by this study was measured in identical conditions (b). **c.** Western-blot showing the expression of hRIPK3 and its variants in transduced HeLa cells. GAPDH is shown as loading control.

**Supplementary Figure 11.**
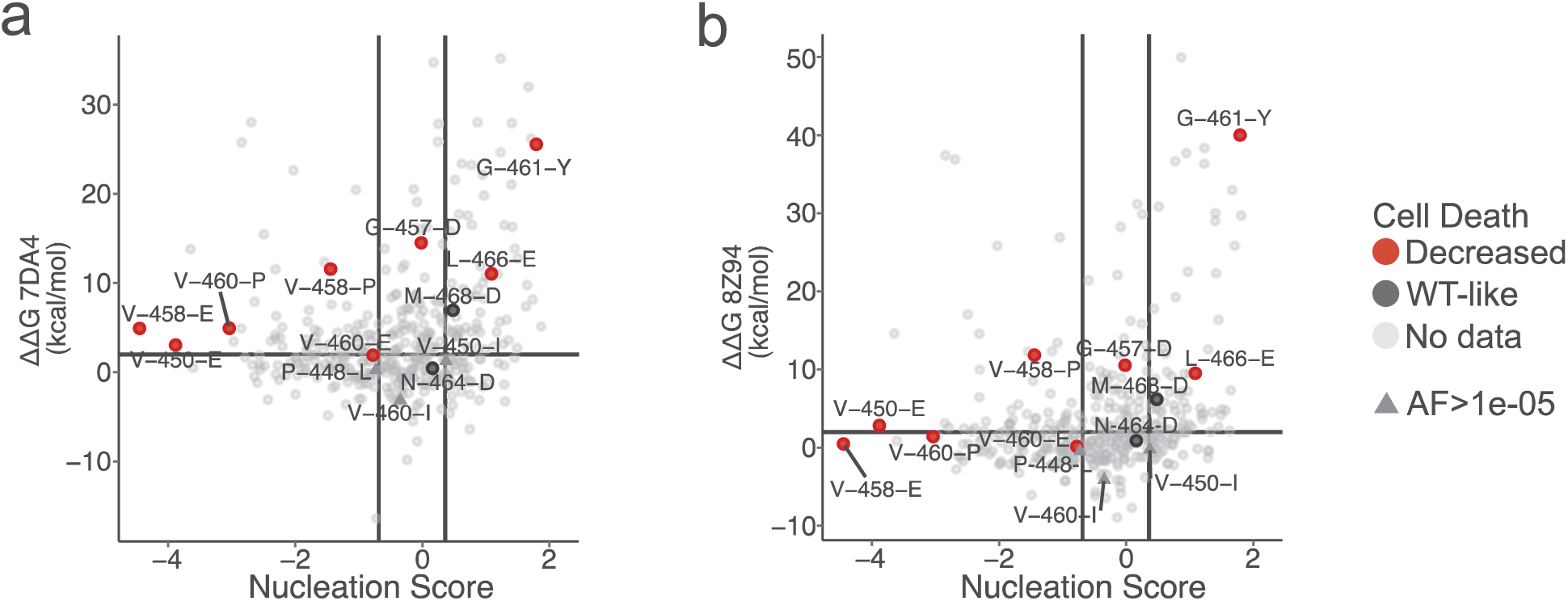
RIPK3 function requires an optimal range of amyloid nucleation and fibril stability. **a** and **b.** Scatter plot of hRIPK3 mutations ΔΔG in 7DA4^20^ structure (a) or 8Z94^22^ structure (b) and nucleation scores. Variants previously studied in the literature, more frequent in the human population (Gnomad Allele Frequency>1e-05) and functionally characterized in this work are highlighted in the scatter plot.

## References

1. Vince, J. E., Davidson, N. M. & Tanzer, M. C. Necroptotic cell death consequences and disease relevance. Nat. Immunol. 26, 1863–1876 (2025).

2. Sun, X., Yin, J., Starovasnik, M. A., Fairbrother, W. J. & Dixit, V. M. Identification of a Novel Homotypic Interaction Motif Required for the Phosphorylation of Receptor-interacting Protein (RIP) by RIP3. Journal of Biological Chemistry 277, 9505–9511 (2002).

3. Cho, Y. et al. Phosphorylation-Driven Assembly of the RIP1-RIP3 Complex Regulates Programmed Necrosis and Virus-Induced Inflammation. Cell 137, 1112–1123 (2009).

4. Li, J. et al. The RIP1/RIP3 Necrosome Forms a Functional Amyloid Signaling Complex Required for Programmed Necrosis. Cell 150, 339–350 (2012).

5. Cuchet-Lourenço, D. et al. Biallelic *RIPK1* mutations in humans cause severe immunodeficiency, arthritis, and intestinal inflammation. Science (1979). 361, 810–813 (2018).

6. Tao, P. et al. A dominant autoinflammatory disease caused by non-cleavable variants of RIPK1. Nature 577, 109–114 (2020).

7. Lalaoui, N. et al. Mutations that prevent caspase cleavage of RIPK1 cause autoinflammatory disease. Nature 577, 103–108 (2020).

8. Liu, Z. et al. Encephalitis and poor neuronal death–mediated control of herpes simplex virus in human inherited RIPK3 deficiency. Sci. Immunol. 8, (2023).

9. Demarco, B., Chen, K. W. & Broz, P. Cross talk between intracellular pathogens and cell death. Immunol. Rev. 297, 174–193 (2020).

10. Verdonck, S., Nemegeer, J., Vandenabeele, P. & Maelfait, J. Viral manipulation of host cell necroptosis and pyroptosis. Trends Microbiol. 30, 593–605 (2022).

11. Pham, C. L. et al. Viral M45 and necroptosis-associated proteins form heteromeric amyloid assemblies. EMBO Rep. 20, (2019).

12. Shanmugam, N. et al. Herpes simplex virus encoded ICP6 protein forms functional amyloid assemblies with necroptosis-associated host proteins. Biophys. Chem. 269, 106524 (2021).

13. Steain, M. et al. Varicella zoster virus encodes a viral decoy RHIM to inhibit cell death. PLoS Pathog. 16, e1008473 (2020).

14. Riebeling, T., Kunzendorf, U. & Krautwald, S. The role of RHIM in necroptosis. Biochem. Soc. Trans. 50, 1197–1205 (2022).

15. Liu, Q., Liu, C., He, Q., Wang, L. & Song, L. The involvement of CgRHIM-containing protein in regulating haemocyte apoptosis after high temperature stress in Pacific oyster Crassostrea gigas. Dev. Comp. Immunol. 159, 105226 (2024).

16. Fay, E. J., Isterabadi, K., Rezanka, C. M., Le, J. & Daugherty, M. D. Evolutionary and functional analyses reveal a role for the RHIM in tuning RIPK3 activity across vertebrates. Elife 13, (2025).

17. Dyrka, W. et al. Identification of NLR-associated Amyloid Signaling Motifs in Bacterial Genomes. J. Mol. Biol. 432, 6005–6027 (2020).

18. Liu, J. et al. The structure of mouse RIPK1 RHIM-containing domain as a homo-amyloid and in RIPK1/RIPK3 complex. Nat. Commun. 15, 6975 (2024).

19. Wu, X. et al. The amyloid structure of mouse RIPK3 (receptor interacting protein kinase 3) in cell necroptosis. Nat. Commun. 12, 1627 (2021).

20. Wu, X. et al. The structure of a minimum amyloid fibril core formed by necroptosis-mediating RHIM of human RIPK3. Proceedings of the National Academy of Sciences 118, (2021).

21. Polonio, P. et al. Structural basis for amyloid fibril assembly by the master cell-signaling regulator receptor-interacting protein kinase 1. Nat. Commun. 16, 9607 (2025).

22. Ma, Y. et al. Intercellular propagation of RIPK1/RIPK3 amyloid fibrils. Proceedings of the National Academy of Sciences 122, (2025).

23. Mompeán, M. et al. The Structure of the Necrosome RIPK1-RIPK3 Core, a Human Hetero-Amyloid Signaling Complex. Cell 173, 1244–1253.e10 (2018).

24. Yang, D. et al. ZBP1 mediates interferon-induced necroptosis. Cell. Mol. Immunol. 17, 356–368 (2020).

25. Chen, X. et al. Mosaic composition of RIP1–RIP3 signalling hub and its role in regulating cell death. Nat. Cell Biol. 24, 471–482 (2022).

26. Chandramowlishwaran, P. et al. Mammalian amyloidogenic proteins promote prion nucleation in yeast. Journal of Biological Chemistry 293, 3436–3450 (2018).

27. Seuma, M., Faure, A. J., Badia, M., Lehner, B. & Bolognesi, B. The genetic landscape for amyloid beta fibril nucleation accurately discriminates familial Alzheimer’s disease mutations. Elife 10, (2021).

28. Cheng, J. et al. Accurate proteome-wide missense variant effect prediction with AlphaMissense. Science (1979). 381, (2023).

29. Brandes, N., Goldman, G., Wang, C. H., Ye, C. J. & Ntranos, V. Genome-wide prediction of disease variant effects with a deep protein language model. Nat. Genet. 55, 1512–1522 (2023).

30. Laine, E., Karami, Y. & Carbone, A. GEMME: A Simple and Fast Global Epistatic Model Predicting Mutational Effects. Mol. Biol. Evol. 36, 2604–2619 (2019).

31. Guerois, R., Nielsen, J. E. & Serrano, L. Predicting Changes in the Stability of Proteins and Protein Complexes: A Study of More Than 1000 Mutations. J. Mol. Biol. 320, 369–387 (2002).

32. Hu, H. et al. RIP3-mediated necroptosis is regulated by inter-filament assembly of RIP homotypic interaction motif. Cell Death Differ. 28, 251–266 (2021).

33. Zhang, H. et al. Crucial Roles of the RIP Homotypic Interaction Motifs of RIPK3 in RIPK1-Dependent Cell Death and Lymphoproliferative Disease. Cell Rep. 31, 107650 (2020).

34. Wu, E. et al. HSPA8 acts as an amyloidase to suppress necroptosis by inhibiting and reversing functional amyloid formation. Cell Res. 33, 851–866 (2023).

35. Sun, L. et al. Mixed Lineage Kinase Domain-like Protein Mediates Necrosis Signaling Downstream of RIP3 Kinase. Cell 148, 213–227 (2012).

36. Sun, K. Y. et al. A deep catalogue of protein-coding variation in 983,578 individuals. Nature 631, 583–592 (2024).

37. Chen, S. et al. A genomic mutational constraint map using variation in 76,156 human genomes. Nature 625, 92–100 (2024).

38. Faure, A. J., Schmiedel, J. M., Baeza-Centurion, P. & Lehner, B. DiMSum: an error model and pipeline for analyzing deep mutational scanning data and diagnosing common experimental pathologies. Genome Biol. 21, 207 (2020).

39. Larsen, J. A. et al. The mechanism of amyloid fibril growth from Φ-value analysis. Nat. Chem. 17, 403–411 (2025).

40. Humphrey, W., Dalke, A. & Schulten, K. VMD: Visual molecular dynamics. J. Mol. Graph. 14, 33–38 (1996).

41. Pettersen, E. F. et al. UCSF Chimera—A visualization system for exploratory research and analysis. J. Comput. Chem. 25, 1605–1612 (2004).

42. Tordai, H. et al. Analysis of AlphaMissense data in different protein groups and structural context. Sci. Data 11, 495 (2024).

43. Sievers, F. et al. Fast, scalable generation of high-quality protein multiple sequence alignments using Clustal Omega. Mol. Syst. Biol. 7, (2011).

